# How spatial attention affects the decision process: looking through the lens of Bayesian hierarchical diffusion model & EEG analysis

**DOI:** 10.1101/2021.05.12.443763

**Authors:** Amin Ghaderi-Kangavari, Kourosh Parand, Reza Ebrahimpour, Michael D. Nunez, Jamal Amani Rad

## Abstract

Model-based cognitive neuroscience elucidates the cognitive processes and neurophysiological oscillations that lead to behavioral performance in cognitive tasks (e.g., response times and accuracy). In this paper we explore the underlying latent process of spatial prioritization in perceptual decision processes, based on one well-known sequential sampling model (SSM), the drift-diffusion model (DDM), and subsequent nested model comparison. Neural components of spatial attention which contributed to the latent process and behavioral performance in a visual face-car perceptual decision were detected based on both time-frequency decomposition and event-related potential (ERP) analysis. For estimating DDM parameters (i.e. the drift rate, the boundary separation, and the non-decision time), a Bayesian hierarchical approach is considered, which allows inferences to be performed simultaneously on the group and individual level. Our cognitive modeling analysis revealed that spatial attention changed the non-decision time parameter across experimental conditions, such that a model with a changing non-decision time parameter provides a better fit to the data than other model parameters, quantified using the deviance information criterion (DIC) score and R-squared. Using multiple regression analysis on the contralateral minus neutral N2 sub-component (N2nc) at central electrodes, it can be concluded that poststimulus N2nc can predict mean response times (RTs) and non-decision time parameters related to spatial prioritization. However the contralateral minus neutral alpha power (Anc) at parieto-occipital electrodes can only predict the mean RTs and not the non-decision time relating to spatial prioritization. It was also found that the difference of contralateral minus neutral neural oscillations were more reflective of the modulation of the top-down spatial attention in comparison to the difference of ipsilateral minus neutral neural oscillations. These results suggest that individual differences in spatial attention are encoded by contralateral (and not ipsilateral) N2 oscillations and non-decision times. This work highlights how model-based Cognitive Neuroscience can further reveal the role of EEG in spatial attention during perceptual decision making.

## Introduction

Decision-making is a high-level cognitive process whereby the decision-maker makes decisions based on the available evidence, expected value, and the possible outcomes (^1^). This process, the decision, is thought to operate in a flexible time frame and is influenced by the real-time demands of the action itself, instead of in the form of immediate evidence acquisition (^2, 3^). The most frequent type of decision that we routinely make in different situations is perceptual decision making, a process that involves the processing of information, the accumulation of evidence, and motor response (^4^).

Spatial attention is also known to play a key role in the perceptual decision-making process (^5–8^). Suppose that you are driving to an intersection on a sunny day with the help of traffic lights; it is easy to determine whether to continue driving or stop the vehicle. But if the situation changes and sensory information is disturbed, then response time and accuracy are affected. One example is that when it is rainy and dark (i.e. a decrease in visual coherence) a driver will not effortlessly detect the color and symbol of the traffic light, and thus their ability to stop or accelerate could be affected. A similar, but different effect, is when a driver is moving toward the intersection in rainy and dark weather, and she/he does not know the exact spatial location of the traffic light. The driver could initially struggle to locate the traffic light (i.e. the stimulus) before processing the received information from the light. If the driver is told exactly where the light is (e.g. spatial cueing) before reaching the intersection, the driver could focus their attention to the target and suppress attention to other areas of visual space faster and maybe more accurately (^3, 9–11^). Posner’s (^12^) studies have shown that people can pay close attention to an area of space without moving their eyes to that area. Moreover, one type of spatial attention is goal-driven, suggesting top-down or endogenous attention in situations where we attend to a region of space where the stimulus is present (i.e. valid) or not present (i.e. invalid) (^12^). The top-down cue could facilitate behavioral performance and manipulate neural mechanisms (^13, 14^), of which some aspects are still unknown. In this paper we focus on valid, covert spatial attention to stimuli which were presented by endogenous orienting cues.

One biomarker with a rich history in the study of spatial attention is the alpha band (8-13 Hz) given by frequency and time-frequency compositions of EEG. Top-down spatial attention is thought to modulate the alpha band (8-13 Hz) of down stream areas such as posterior-occipital sites to direct and allocate limited processing resources to attended and unattended loci (^15, 16^). In fact, studies show that increasing alpha band power ipsilaterally (alpha-synchronization) to the unattended location could suppress task-irrelevant space, and the decreasing alpha band power contralaterally (alpha-desynchronization) to the attended location intentionally facilitates upcoming visual processing (^15, 17^). This mechanism of top-down orienting of attention is usually found after post-cue (pre-stimulus) of various tasks such as visual (^18, 19^), tactile (^20^), and auditory (^21^) tasks.

Another class of EEG biomarker that is of interest in spatial attention are the negative deflections in trial-averaged ERPs after visual stimuli occur, in particular the second negative peak (N2) amplitudes. N2s have multiple interpretations and differences in amplitude and latency depending upon the task and location over electrodes. One example is the central contralateral N2 (N2cc) subcomponent in central electrode sites. The N2cc has been shown to have a substantial role in the prediction of distributed finite resources for spatial prioritization (^22^). The N2cc amplitude, which is preparation mechthought to mirror a motor anism, decreases with informative top-down cues (^22^). The N2cc may also reflect an attention-related motor component which could prevent the release of incorrect responses after presentation of stimuli in the ipsilateral field (^23^). Moreover, researchers have reported that the N2cc over central electrodes sites might coincide in time with the posterior contralateral N2 (N2pc), but that the important finding of N2cc is not due to overlap of the N2pc with movement execution-related activity (^22, 24^). It has also been shown that the age-related behavioral was related to N2pc and N2cc subcomponent during visual search task (^23^). The N2pc was found to play a role in visuospatial processing of target stimuli and the control of non-target stimuli, and the N2cc was also correlated to inhibition of spatial response tendencies in the Simon task (^25^). Another similar biomarker, the anterior contralateral N2 subcomponent (N2ac) has been shown to predict sound localization performance and diffusion model parameters using multiple linear regression^5, 26^). In particular it has been shown that N2ac amplitudes modulated accuracies and evidence accumulation rates estimated by drift-diffusion models (DDMs) (^5^). Finally, generalized contralateral (N2c) and ipsilateral (N2i) N2 subcomponent peak amplitudes at electrodes P7/P8 have been found to change as a function of motion strength (^27^), and generalized N2 amplitudes have been shown to track non-decision times in DDMs (^28^).

EEG signals thus contain informative signals encoding spatial attention that describe the allocation of resources to speed up perceptual decision making. However in the literature it is unclear if these EEG signals are contralateral or ipsilateral to the attended stimulus. It could be that some contralateral and ipsilateral EEG biomarkers of spatial attention do not describe anything about spatial attention, and this would not be clear from the literature since most studies calculate EEG biomakers from the difference in contralateral and ipsilateral signals (^5, 25, 26, 29^). Therefore in our study we purposely compared contralateral and ipsilateral EEG signals to a neural condition, in order to better understand how EEG biomarkers describe cognition through direct testing on cognitive variables and behavior.

In the research presented here, we found biomarkers of spatial prioritization in the electroencephalogram (EEG) to predict behavioral performance and DDM parameters in a face-car perceptual decision making task with spatial cueing manipulations. To be more specific, we tested whether the lateralized alpha frequency band at parieto-occipital sites together with the contralateral-neutral (contralateral minus neutral difference curve) and ipsilateral-neutral (ipsilateral minus neutral difference curve) N2 subcomponents at central sites could modulate behavioral performance and parameters of spatial prioritization in perceptual decision making (^5, 24^). The goal of this study was to find the relationship between these two intriguing biomarkers, e.g. the lateralized alpha frequency band amplitudes over parieto-occipital electrode sites and N2-subcomponent amplitudes, with behavioral performance and the non-accumulation time, also called the non-decision time, as estimated from DDMs (^30^). We applied hierarchical Bayesian modeling with a powerful nested model comparisons approach to estimate DDM model parameters and find the model that best described the data (^31^). Then we found out how well individual differences in EEG measures of spatial prioritization described individual differences in task behavior and DDM parameters.

### 0.1 Cognitive modeling of perceptual decisions

The purpose of behavioral modeling is to understand what components or parameters can be extracted from the perceptual decision making process, how different people make correct responses and commit errors, and the time course of these processes (^32^). There is a growing consensus that the evidence accumulates gradually and sequentially to make a decision (^30, 33, 34^). As a result, sequential sampling models have become the most well-known explanation of how the decision-making process works (SSMs; see Stone, 1960 (^35^), Ratcliff, Smith, Brown, and McKoon, 2016 (^36^) and Evans and Wagenmakers, 2020 (^37^) for reviews). These models have been successful in describing the cognitive processes underlying decision making across a wide variety of paradigms, such as studies of optimality (^38–41^), stop signal paradigms (^42^), go/no-go paradigms (^43^), multi-attribute & many alternatives choice (^44–46^), learning strategies (^47–49^), attentional choice (^6–8, 50^), continuous responses (^33, 51^), neural processes (^1^), and so on.

In general, SSMs assume that decisions are made from a noisy process of accumulating evidence where the evidence is gradually accumulated over time until a sufficient amount of evidence for one of the choices reaches a predetermined threshold, at which point a decision is made. DDMs assume particular random processes for accumulating evidence, “Wiener” or Brownian motion-based processes. The reason for this assumption is that evidence accumulation is thought to be a process in which the individual may respond differently to the exact same stimulus on different trials (^36, 52^). DDMs can also be considered as a bridge between behavioral data and cognitive processes because DDMs have cognitive parameters that have been experimentally-supported interpretations (^36, 53^).

Hitherto, various sequential sampling models have been presented by researchers. There are some successful sequential sampling models such as: EZ-diffusion model (^54^), linear ballistic accumulators (^55^), leaky-competing accumulator (^56^), and race diffusion model (^57^), but the drift-diffusion model (DDM) is the most popular model of evidence accumulation (^34^). Similar to the other sequential sampling models, the DDM is based on evidence accumulation until reaching a threshold. In other words, DDM assumes that the information accumulation process starts from a point between two fixed boundaries, the accumulator steps toward the upper/lower boundary, and after the accumulator passes one of these boundaries, the accumulation process stops and the corresponding option is selected (^58^), illustrated in Figure 1. Mathematically, evidence accumulation in the DDM can be described continuously as,

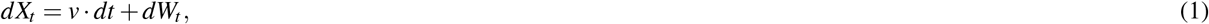

or approximated discretely as,

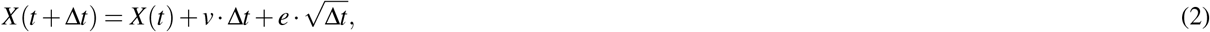

in which *X*_*t*_ is the diffusion state and *dW*_*t*_ denotes the standard Wiener process. Also in discrete form (simulated in Figure 1), *e* is normally distributed noise *N*(0, 1). Traditionally, the DDM consists of four major parameters. The first parameter is the threshold. In DDM the lower boundary is at zero and the upper boundary has distance ‘*a*’ from the lower boundary. Therefore, ‘*a*’ is the threshold parameter, and actually is an index for the speed-accuracy tradeoff. That implies that for higher values of ‘*a*’, the participant accumulates more evidence to make a decision, and thus his/her decisions will be accurate more often, but often slower. On the other hand, lower values of ‘*a*’ imply that the participant often makes the decision faster but will be accurate less often (^39^). The second parameter of the DDM is the drift rate, which is denoted by ‘*v*’. The drift rate is the mean slope of each step of the accumulator toward the boundaries. This parameter encodes the speed of information processing and identifies task difficulty level. Higher values of the drift rate show that the task has no demanding cognitive load and that it is easy to make a decision between the options. When the value of the drift rate tends to zero, this implies the task is demanding and the speed of information processing is slow (^59^). The third parameter is the bias parameter, which denoted by ‘*z*’, that encodes the starting point of the accumulation process. When the starting point is equal to 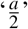, the distance between the starting point and the boundaries are equal and there is no bias to each option. But when ‘*z*‘ is greater or less than 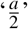, then the participant has a bias to upper or lower boundaries, respectively (^60^). The fourth parameter is the non-accumulation time *T*_*er*_, also called the non-decision time. This parameter is added to the model for the purpose of excluding the encoding and motor time for the response time. Usually, this parameter is denoted by *T*_*er*_ in DDM (^53^). These are the main parameters of the DDM, however often some between-trial variability in parameters is assumed. *s*_*v*_, *s*_*z*_, and *s*_*t*_ are the between trial variability parameters for drift rate, bias, and non-decision time respectively. Usually, normal distributions are assumed for *s*_*v*_ and uniform distributions for *s*_*z*_ and *s*_*t*_ (^30^). High values for the between trial variability parameters indicate that there is some variability in the stimuli across trials or in cognitive states (^61^).

**Figure 1.**
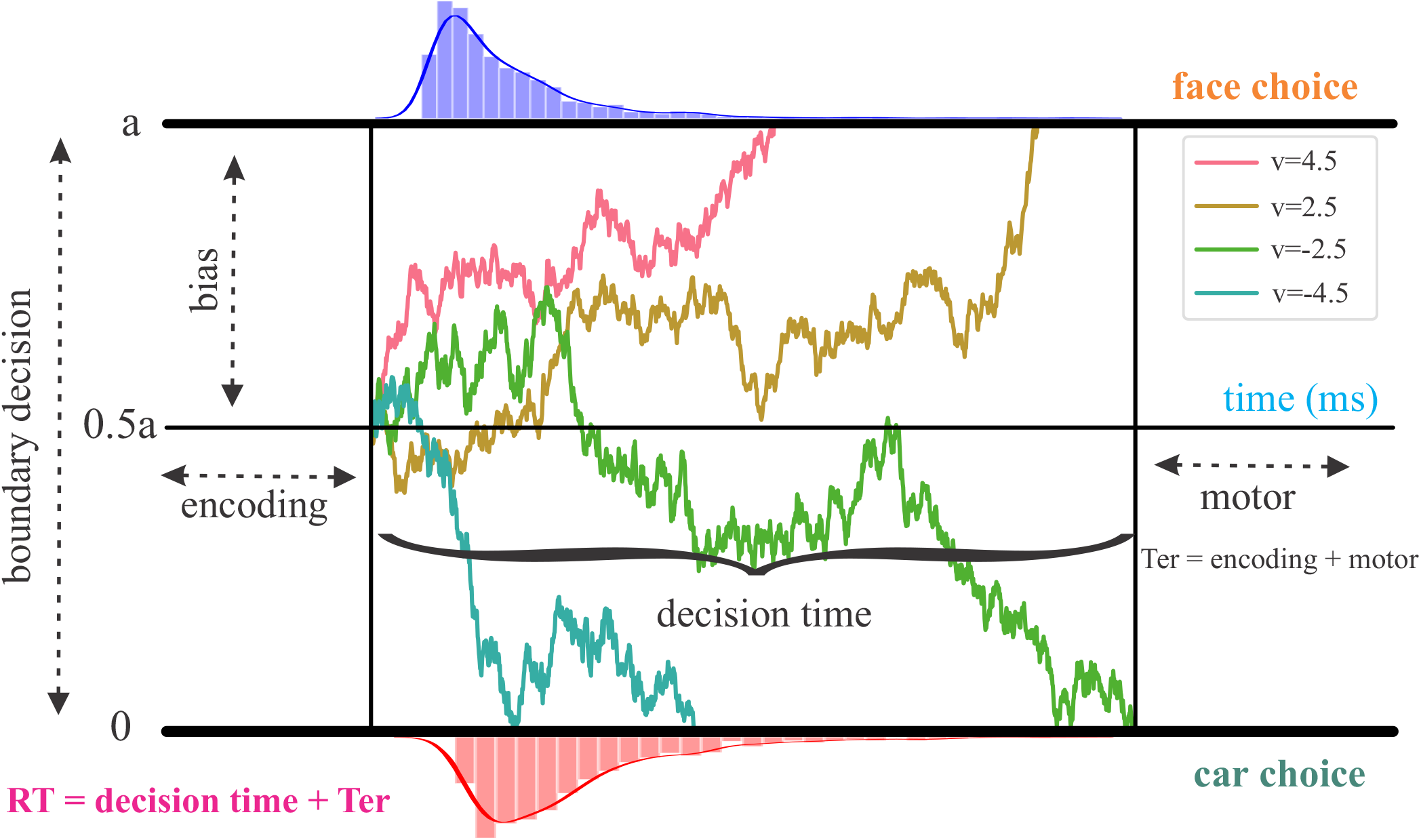
Trajectories of drift-diffusion model for two stimuli with various drift rates. Four trajectories of evidence accumulation are shown on 4 trials with a different drift-rate *v* in each trial. Pink, yellow, green, and blue trajectories were generated with drift rates of 4.5, 2.5, –2.5, and −4.5 respectively. When enough evidence is reached by crossing one of two boundaries for a “face” or “car” choice, a decision is made on that trial. The time course of the trajectories encode the decision time on that trial. The decision time shown in this figure is of the trial given by the green trajectory. “encoding” and “motor” non-accumulation times are assumed to be the two additive components of non-decision time *T*_*er*_. On these four trials there was no initial bias towards a “face” or “car” choice, such that the start point is equal to 0.5*a*.

In summary, we hypothesize that electrophysiological correlates of top-down spatial prioritization (alpha band lateralization and N2 subcomponents) during a visual perceptual decision making task could predict visual performance and the non-decision time parameter of DDMs, reflecting the sensory coding time and response expectation. Our claim is that the cognitive process of spatial attention is encoded by lateralized alpha power in and around parieto-occipital electrode sites, and that spatial attention is also encoded by lateralized N2cc amplitudes in central electrode sites. These measures of spatial attention in EEG should regulate the speed to choose a face and car stimuli, so they should contribute to the non-decision processes and the performance.

## 1 Methods

### 1.1 Experiment overview

In this study, we re-examined the data from an experiment conducted by (^62^) to understand visual coherence and spatial attention effects in perceptual decision-making. In their study, seventeen participants (8 females, mean age was 25.9 years, range 20-33 years, 2 left-handed) from the University of Birmingham completed the experiment, which was approved by the University of Birmingham Ethical Review Committee, and were paid 7.5 per hour with travel expenses reimbursed. Electroencephalograms (EEG) and behavioral data from a perceptual decision making task with spatial cueing were acquired over two data acquisition sessions.

As shown in Figures 2a and 2b, on each experimental trial, in order to manifest the informative and uninformative spatial prioritization, cueing one-way arrows or two-way arrows were shown for a one second period before the stimulus emergence. Then, a visual stimulus of a car or face was displayed on the screen for 200 milliseconds. Participants then pressed a button to respond whether they perceived the car or the face. Participants used their index finger and middle finger of the right hand to respond. This was followed by an inter-trial period between 0 msec and 300 ms. It should be noted that participants were also told to maintain fixated on the fixation point during the entire trial.

**Figure 2.**
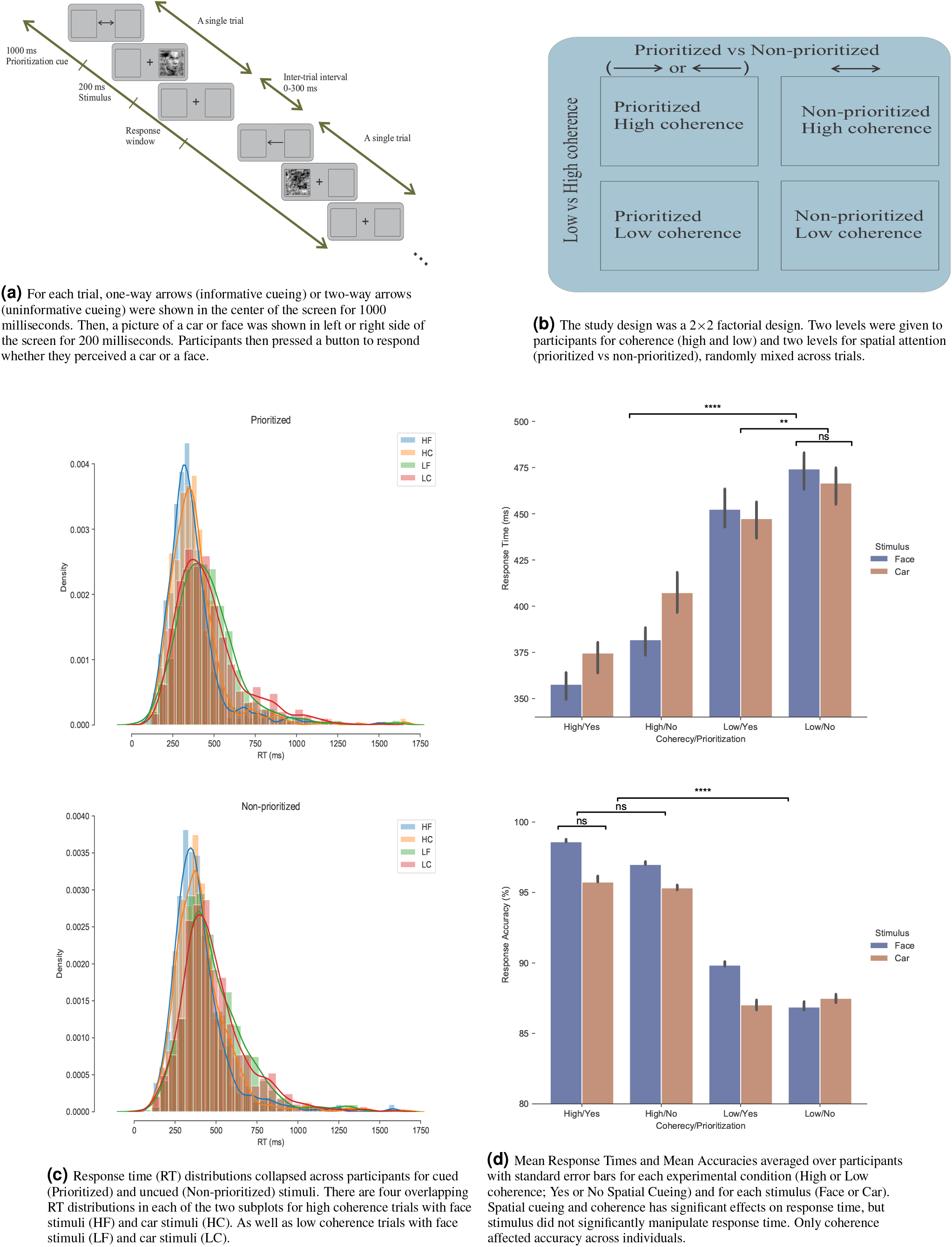
Experimental design and behavioral data. (A-B) Overview of the experiment task and the 2 × 2 study design. (C) Response time (RT) distributions of face and car for all four conditions (two coherences and two spatial cueing conditions). (D) Bar plots and error bars displayed the grand average and ± 1 standard error of the mean (SEM) response time across independent variables. Asterisks indicate significant p value as **p<0.01 and ****p<0.0001, and ‘np’ indicates non-significant p value.)

The experimental manipulation of coherence had two levels: “low” and “high”, and experimental manipulation of spatial cueing had also two levels: “yes” and “no”, such that a leftward or rightward facing arrow was presented in half of the trials (“yes”) and a double sided arrow was presented in half of the trials (“no”). These arrows indicated to participants that they should attend to one side (informative cueing) of space or both sides of space (uninformative cueing) covertly. All trials from each experimental condition and experimental stimuli (i.e. face or house) were randomly presented to participants. For more details about experimental design & stimuli, interested readers can refer to the main report written on the data by Georgie et al. (^62^).

Following (^62^), 36 trials of both EEG and behavioral data were acquired for each of the experimental conditions (coherence and spatial attention) and two stimulus types (face or house). The EEG was collected with a sampling rate of 5000 Hz samples using 64 EEG sensors, which consisted of 62 scalp electrodes via the 10-20 system and two extra sensors. One extra sensor was placed approximately 2 cm under the left collarbone in an effort to identify electrocardiogram (ECG) signals, and the other sensor was placed under the left eye to aid in correcting eye-blinking artefacts (EOG).

In our study, three participants were excluded because their EEG data were too contaminated with artifact, which left fourteen participants in our analysis. Also, in order to perform statistical analysis and model fitting, we removed outliers in the behavioral data. We used an interquartile range (IQR), with IQR = Q3 - Q1 as the distance from the first quartile to the third quartile of response times. To compute the interquartile range for the data, any response time smaller than Q1 - 3 × IQR and greater than Q3 + 3 × IQR were removed as outliers. Consequently, response time and accuracy are submitted to parametric 2 × 2 × 2 repeated-measure analysis of variance (ANOVA) statistical test.

### 1.2 EEG signal analysis

The EEG preprocessing and analysis were implemented by the MNE-python package (^63^). The preprocessing steps we performed can be summarized as follows: the continuous EEG data were down-sampled to 256 samples, the EEG data were band-pass filtered to the range of 1 Hz - 30 Hz, EEG was re-referencing to the common average, visual inspection of the EEG was performed to remove trials with high noise amplitude or abnormal activity, signals containing too much noise were interpolated, trials with amplitudes outside the range of -100 mV to +100 mV were removed, EEG data was split into epochs from -100 msec (baseline) time-locked to the cue’s onset to 600 msec after stimulus appearance (totalling 1700 ms), Independent Component Analysis (ICA) was run to manually remove EEG data irrelevant to the task (e.g. eye movement, head motion and muscular activity), Independent Components (ICs) were removed automatically that best matched EOG sensors or ECG sensors (such that these ICs reflected eye blinks and heart rhythm artifacts respectively) using the MNE-package (^63^), and finally the EEG data was converted back into sensor space from IC space. The EEG preprocessing with MNE-package implementation code in python named ’MNE-preprocessing’ is readily accessible at https://github.com/AGhaderi/MNE-Preprocessing.

In order to explore the spectral content of the data, we employed a Morlet wavelet transform to the EEG in Python. The Morlet wavelet is an efficient time-frequency decomposition over every trial. We used this time-frequency approach in order to appropriately understand the properties of the non-stationary EEG signal, and to keep the random phase information before averaging across trials (in comparison to the event-related potential analysis discussed below, which only extracts phase-locked information across trials). We changed the parameter of “number of cycles”, which controls the trade-off between temporal precision and frequency precision in this algorithm, by increasing by 0.5 cycles per frequency, starting at 4 Hz, in steps of 1 Hz. That is, we used 3-cycle wavelets at the lowest frequency (4 Hz), 3.5-cycle wavelets at the next lowest frequency (5 Hz), and so on until we reached 16-cycle wavelets at the highest frequency (30 Hz). After performing this single-trial Wavelet decomposition from 4Hz to 30Hz, the time-frequency decomposition per trial was averaged across trials for each participant. We used this time-frequency average across trials to extract the alpha band from 8 to 13 Hz per participant by averaging the power from 8 to 13 Hz.

We also employed an event-related potential (ERP) analysis. This analysis better captures the phase-locked information across trials (as opposed to the time-frequency analysis discussed previously). Because EEG data routinely contains both signal from real brain sources but also from biological or non-biological artifact and noise even after rigorous artifact correction (^29, 64^), a phase-locked analysis is useful to better average-out or eliminate outlier artifact. Therefore, we used an ERP approach to detect cognitive components across experimental conditions and for each participant. We used a baseline of 100 msec before the cue onset in order to calculate the N2 amplitudes after the stimulus appearance.

Both the N2 ERP subcomponent waveforms and alpha band frequency amplitudes were viewed by eye before being used in regression models. We calculated the mean contralateral and ipsilateral (informative) and neutral evoked potentials (uninformative) at C1/2/3/4 central electrodes, and also the mean ipsilateral and contralateral and neutral power across time in the alpha frequency band (8–13 Hz) at parieto-occipital electrodes PO3/4/7/8 (^22–24^). We calculated the contralateral responses as the average of ERP oscillations in left hemispheric sensors during right-arrow cue trials and in right hemispheric sensors during left-arrow cue trials. Similarly we calculated the ipsilateral responses as the average ERP oscillations at left hemispheric sensors in left-arrow cue trials and right hemispheric sensors in right-arrow trials. We also calculated the neutral responses as the average of ERP oscillations at both left and right hemispheric sites in two-way arrow trials.

Many spatial attention studies have focused on pre-stimulus components and alpha oscillations (^13, 65^), and more specifically previous researchers have focused on pre-stimulus alpha oscillations as the anticipatory indexes for allocating of spatial prioritization (^66^). However we concentrated on post-stimulus components during perceptual decision making task to target the actual deployment of spatial attention (^5^). In order to measure the N2 subcomponent amplitudes, we computed the dependent t-test statistic over contralateral (at C1 & C3 or C2 & C4 in informative cue conditions) minus neutral (at C1, C2, C3, and C4 in uninformative cue conditions) amplitude (N2nc) and ipsilateral minus neutral amplitude (N2ni) at a time window within 150-500 msec after stimulus onset (1150-1500 msec after cue onset). This resulted in 90 statistical t-tests, with a test for each time point in the 350 msec window with a 256 sample rate, so we used a False Discovery Rate (FDR) correction procedure that gave us a different standard for significance to correct the t-tests (^29^). The measurement window of N2nc and N2ni amplitudes was then based on significant time samples by FDR correction. We then used the variables of {N2nc}, and {N2ni} were used as regressors in the multiple regression model described later.

To assess alpha lateralization, two contralateral and ipsilateral alpha band indices were used across both hemispheres as follows:

*Anc* = contralateral alpha power (at PO3 & PO7 or PO4 & PO8 in informative cue conditions) minus neutral alpha power (at PO3, PO4, PO7, and PO8 in uninformative cue conditions)

*Ani* = ipsilateral alpha power (at PO3 & PO7 or PO4 & PO8 in informative cue conditions) minus neutral alpha power (at PO3, PO4, PO7, and PO8 in uninformative cue conditions)

When the Anc index is negative, the contralateral alpha power is lower than the neutral alpha power at parieto-occipital sensors and vice versa. In contrast, when the Ani index is positive, the ipsilateral alpha power is higher than the neutral alpha power and vice versa. To determine the measurement window of alpha power in the grand-averaged Anc and Ani indices, we used the 50% Fractional Area Latency (FAL). The FAL estimates the midpoint latency of a component signal (^5, 29^). We calculated 50% FAL contingent on a wide time window, from -100 to 600 msec after stimulus onset (i.e. 900-1600 msec after cue onset). We then used a 200 msec time window around the FAL to extract relevant Anc and Ani. The variables of {Anc}, and {Ani} were then used as regressors in the multiple regression model in addition to the {N2nc}, and {N2ni} regressors mentioned previously.

### 1.3 Cognitive modeling and model comparison

We used a hierarchical Bayesian estimation approach in order to explore the latent cognitive processes underlying perceptual decisions during spatial prioritization, and to link those latent cognitive processes to neural mechanisms. We assumed hierarchical models where the participant-specific parameter values are randomly sampled from a group-level distribution. And then we used a hierarchical Bayesian approach such that the marginal distributions of the parameters are estimated simultaneously at the group level and the individual level (^67^) for each model. We combined this hierarchical approach with the diffusion model in a stable estimation of parameters (i.e. drift rate, the boundary separation, and the non-decision time) at both individual and group levels. Specifically we fit models using the HDDM package in Python (^68^), which generates full posterior distributions of the model parameters at the individual and group level via Markov-chain Monte Carlo sampling (MCMC). We used the HDDM package because is flexible and user-friendly. We also applied hierarchical Bayesian estimation because it returns stable results and appropriate parameter estimation in the form of posterior distributions for each parameter included in the model, even when less data (i.e., experimental trials) are available compared to other implementations such as traditional maximum likelihood estimation for individual participants (for details about the model see^68, 69^).

The HDDM package offers different model parameterizations depending on the experimental factors, e.g., one could model different drift rates for all conditions or for all groups or any combination of them^70^. Different criteria such as the deviance information criterion (DIC) can be used for a comparison across these models. In the full DDM, there are seven parameters which could be categorized into three parts: the decision process measures (boundary decision, mean starting point, and mean drift rate), the non-decision process measure (non-decision time), and the inter-trial variability measures (drift rate variability *s*_*v*_, starting point variability *s*_*z*_, and non-decision time variability *s*_*t*_) (^36, 52^). Based on the strong evidence previously presented in various studies to investigate the effects of coherence and stimulus on the decision-making process (^4, 31, 71^), we first tested cognitive models whose drift rate parameters are dependent both on the coherence condition (high or low) and stimulus (face or car).

In order to test our hypothesis that spatial attention regulated resources of decision making, we compared five possible nested models (^31^). First, the spatial cueing (i.e. directional arrows presented to participants) may not capture or shift any resources or decision parameters, so this model has no parametric dependence on spatial prioritization. We labeled this model *model*_*p*_, in which only the drift rate parameter depends on coherence and stimulus (face or house). Second, spatial cueing could regulate the rate of accumulated evidence (the drift rate *v*), labeled *model*_*v*_. Third, non-decision time (*T*_*er*_) could vary with spatial cueing, labeled *model*_*t*_. Four, spatial cueing could manipulate the mean bias (starting point *z*), labeled *model*_*z*_. Five, spatial cueing might shifts the decision boundary (*a*), called *model*_*a*_. Note that in each model the drift rate *v* was free to vary with the level of the coherence (high or low), in line with related findings about stimulus strength in perceptual tasks (^4, 72^). Moreover, inter-trial variability in non-decision time ’st’ was a free parameter in all models on the group level, whereas other inter-trial variables were not estimated. Note that there are two ways to implement DDMs: an accuracy-based approach in which correct responses are coded 1 and incorrect responses 0, or a stimulus-based approach in which the response choice, such as face or car, are modeled. In this work, the stimulus-based approach is employed in which “face” responses are coded 1 and “car” responses are coded 0.

For each model, Markov Chain Monte Carlo simulations (^73^) were used to generate 100,000 samples from the joint posterior distribution of parameters using the HDDM package (^68^), from which the initial 1000 samples were discarded as a burn-in phase to minimize the effect of initial values on the posterior inference and converge on stable distributions (^68^). The convergence of the Markov chains was assessed through visual inspection as well as by calculating the Gelman-Rubin statistic (^74^), R-hat, to ensure that the models had properly converged, such that R-hat compares between-chain and within-chain variance.

Both R-squared (*R*^2^) and the deviance information criterion (DIC) of each model were used to evaluate each model’s goodness-of-fit while accounting for model complexity (i.e., number of free parameters), with lower DIC values indicating better model fit^75^. Note that DIC have been previously used to choose the best diffusion model on group level (^18, 68^). DIC score is reported for group level of DDMs. *R*^2^ was generating from simulated RT and choice data from the group level DDM parameters across conditions by comparing generated mean RTs and accuracies with observed group mean RTs and accuracies.

To further evaluate the best fitting model, we ran posterior predictive checks by averaging 500 simulations generated from the model’s posterior to confirm it could reliably reproduce patterns in the observed data (^68^), see Supplementary Material. The hDDM implementation code and simulation scripts for all current models are readily available at https://github.com/AGhaderi/hDDM_attention.

### 1.4 Multiple regression model

In this section, we explored whether both pairs of explanatory (independent) variables of {N2nc, Anc}, and {N2ni, Ani} could predict spatial prioritization in the best-fitting model (*model*_*t*_) described in the previous section, as well as behavioral performance (RT and accuracy). Multiple regressions using the Ordinary Least Squares (OLS) method were fit using a statistics module (statsmodels) in Python, with the following formula:

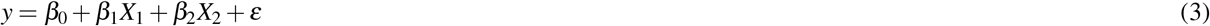

Two regression model types were fit for each pair of regressors. In one model, *X*_1_ was *N*2*nc* and *X*_2_ was *Anc*, and in the other model *X*_1_ was *N*2*ni*, and *X*_2_ was *Ani*. We allowed the dependent variable *y* to be either a direct estimate of behavioral performance (RT or accuracy) or an estimated non-decision time parameter from *model*_*t*_, the best fitting DDM, per participant. In order to investigate the mechanism of spatial attention based on electrophysiological components in perceptual decision making, we applied the separate multiple regression models with prioritization (informative spatial cueing) minus non-prioritization condition (uninformative spatial cueing) dependent variables. That is, the dependent variables of these regressions were calculated by subtracting the non-decision time parameter (or mean RT or accuracy) of the informative spatial cueing condition from the non-decision time parameter (or mean RT or accuracy) of the uninformative spatial. The non-decision time parameters were estimated using mean posterior distribution for each participant from the best fitting model, *model*_*t*_. The model regressors were the pairs of explanatory (independent) variables of {N2nc, Anc}, and {N2ni, Ani} previously mentioned. This resulted in six multiple regressions being run. Each multiple regression model was used to assess the relationship between electrophysiological correlates and latent processing of spatial prioritization resulting from modeling parameters with p-values found for each beta coefficient.

We also calculated a Bayes factor (BF) for each model. BFs, which range from 0 to ∞, quantifies the relative evidence for one model over an another model (in our case, the null model with only an intercept/mean term). For example, if BF_10_ = 5, it means that the evidence is 5 times more probable under the alternative model than under a null model. When the BF equals 1, it indicates that both model models (i.e., the null and the alternative model) have equal probability. Moreover, a BF_10_ > 1 gives the evidence in favor of the alternative model instead of the null model. If the BF_10_ > 3 or > 10, it shows moderate and strong evidence for the alternative model, respectively. We denote the BF with a subscript, BF_10_, to clarify that this BF gives the probability in favor of the alternative model over the null model (^76^). We calculated the BF10 in R using the BayesFactor package (^77^).

In addition, *R*^2^ is reported as a goodness-of-fit statistic that summarizes the relationship between the observed and predicted values for each linear model. Component-Component plus Residual (CCPR) or the partial Residual plot (e.g. graphics.plot_ccpr function from statsmodels module in Python) is a worthwhile metric that can be utilized to assess the impact of a regressor on the outcome variable (y) by taking into account the effects of the other independent variables. The CCPR plot shows the relationship of *β*_*i*_*X*_*i*_ + *ε* versus *X*_*i*_. The CCPR plot also provides the (*β*_*i*_*X*_*i*_, *X*_*i*_) line, so that one can see whether the relationship between the variation in a regressor on response variable seems to be linear or not.

## 2 Results

### 2.1 Behavioral result

Both coherence and spatial cueing were distinct experimental variables, which could have changed the quality of sensory information and the top-down attention to a specific visual field respectively. However, in order to understand the true effects of the experimental manipulation, the behavioral performance was analyzed using three-way repeated measure ANOVA. For the ANOVA of mean response times for each participant, this ANOVA found no significant main effects of stimulus (F(2,13) =0.78, p = 0.39), but revealed significant main effects of spatial cueing (F(2,13) = 11.83, p = 0.0044 < 0.01) and phase coherence (F(2,13) = 38.28, p < 0.0001) (Fig. 2d). Specifically, the easier phase coherence condition and informative spatial cueing condition led to faster response times. Interaction between stimulus and coherence was also significant (F(2,13) = 8.54, p = 0.0119 < 0.05), but the other interactions between variables were not statistically significant, i.e. interaction of spatial cueing and phase coherence (F(2,13) = 0.54, p = 0.47) and interaction of spatial cueing and stimulus (F(2,13) = 0.13, p = 0.71). Because of the significant interaction between coherence and stimulus, it was worthwhile to create DDMs describing response choices (e.g. face or house), and not just DDMs describing trial-by-trial accuracy (e.g. correct or incorrect), so that these effects can be observed with the drift rate. A three-way repeated measure ANOVA was also fit to mean accuracy for each participant. The only significant main effect (F(2,13) = 41.83, p < 0.0001) (Fig. 2d) was that of coherence. The main effects of stimulus (F(2,13) = 0.84, p = 0.37) and spatial cueing (F(2,13) = 2.12, p = 0.17) were not significant. Similarly the interactions of spatial cueing and phase coherence (F(2,13) = 0.05,p = 0.81), of spatial cueing and stimulus (F(2,13) = 2.9, p = 0.11) and of coherence and stimulus (F(2,13)= 0.14, p = 0.71) were not significant.

### 2.2 Cognitive modeling

To explore the underlying mechanisms of spatial attention in perceptual decision making, we fit five hierarchical drift-diffusion models (HDDMs) previously mentioned. The convergence diagnostics of the best fitting model, *model*_*t*_, are reported in the Supplementary Material of the article. MCMC sampling traces of the group posterior distributions showed converged chains during model fitting (see Figures 8, 9 and 10). Furthermore, the convergence of the Markov chains was assessed by calculating the R-hat Gelman-Rubin statistic (^74^). It should be noted that *model*_*t*_ showed superior convergence and the same stationary distribution with four chains, based on R-hat values under 1.0001 for the parameters (see Supplementary Material). Also, we compared posterior predictions from each model with the observed data. There was near agreement between the observed data and the model predictions across conditions in all conditions, and the model is able to capture the overall shape of response times across the four different conditions (see Figure 11). Furthermore, the result of model comparison analysis reveals that model_*t*_ provided the best model fit, in that it had the smallest DIC and largest *R*^2^. This suggests that the non-decision time parameter (*T*_*er*_) is manipulated by spatial cueing (see Table 3), more so than other than other parameters.

We then utilized the results of *model*_*t*_ to examine our hypotheses and identify the electrophysiological mechanisms of informative (contralateral and ipsilateral) and uninformative (neutral) prioritization under decision-making. In order to assess parameters of the winning *model*_*t*_ across conditions, we used a dependent Student’s t-test on non-decision time and two-way repeated measure ANOVA on the drift-rate parameter. Mean non-decision processes of spatial cueing and non-spatial cueing were significantly different (t =3.34, p = 0.0053 < 0.01), see Fig. 3a. That is, we found that non-decision processes take less time when people receive an informative spatial cue than when they receive an ambiguous (uninformative) spatial cue. In addition, the main effect of coherence on drift rate was significant (F(1,13) = 75.11, p < 0.001), while the stimulus had no effect on drift rate (F(1,13) = 0.3, p = 0.87), see Fig. 3b. There were also no significant interaction between stimulus and coherence on drift rate (F(1,13) = 0.64, p = 0.43). Box-plots of non-decision time *T*_*er*_ and drift rate parameters in different conditions are reported in Figs. 3c and 3d, respectively.

**Figure 3.**
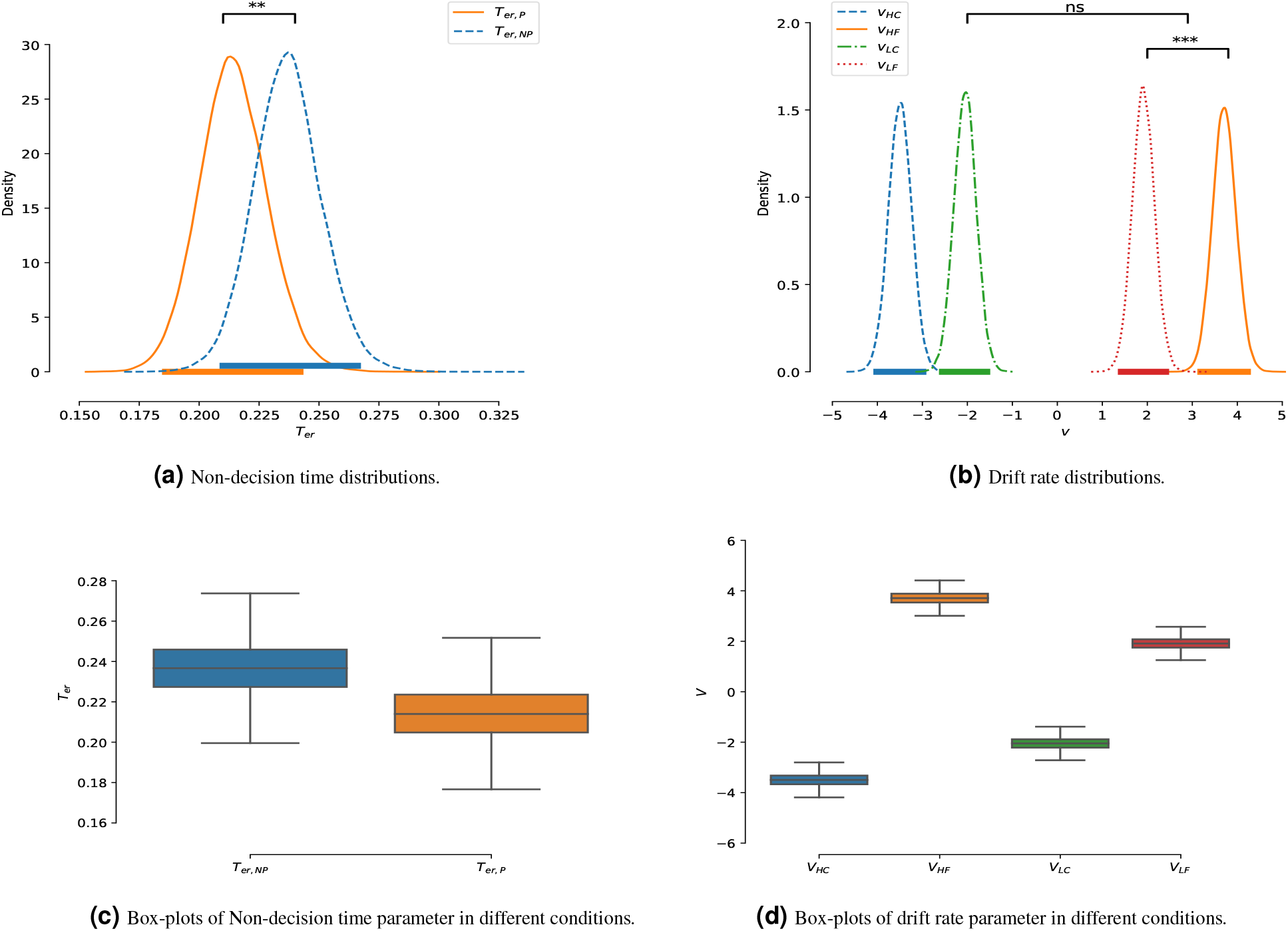
Posterior distribution of the group parameter non-decision time and drift rate from the model_*t*_. (a), Non-decision was found to be different across the two types of cueing experimental conditions (t = 3.34, p = 0.0053 < 0.01). (b) The main effect of coherence on drift rate was significant (F(1,13) = 75.11, p < 0.001), but the main effect of stimulus on drift rate was not significant (F(1,13) = 0.3, P = 0.87). There was also no significant of interaction between stimulus and coherence of drift rate (F(1,13) = 0.64, P = 0.43). Each legend item refers to a different posterior distribution from *model*_*t*_ : *T*_*er,P*_: non-decision time of prioritization (informative spatial cueing), *T*_*er,NP*_: non-decision time of non-prioritization (uninformative spatial cueing), *v*_*HC*_: drift rate of high coherence and car stimulus, *v*_*HF*_ : drift rate of high coherence and face stimulus, *v*_*LC*_: drift rate of low coherence and car stimulus, *v*_*LF*_ : drift rate of low coherence and face stimulus. Asterisks indicate significant p value as **p<0.01 and ***p<0.001, and ’np’ indicates non-significant p value. Thick horizontal lines are 95% of Highest Posterior Density (HPD).

#### Correlation of the DDM and conventional outcome Parameters

To understand which characteristics of behavioral outcome are indicative of cognitive parameters, the correlations were found across participants between mean posterior values of non-decision time from model_*t*_ and mean response times across trials. Matrix correlations are visualized in Figure 4. *RT*_*er,P−NP*_ (response time of prioritization minus response time of non-prioritization) and *T*_*er,P−NP*_ (non-decision time of prioritization minus non-decision time of non-prioritization) is highly positively correlated ‘0.47’, indicating that larger *T*_*er,P−NP*_ in the decision process is associated with longer responses *RT*_*P−NP*_. Therefore, this indicates that RT changes relating to spatial attention are influenced by non-decision time.

**Figure 4.**
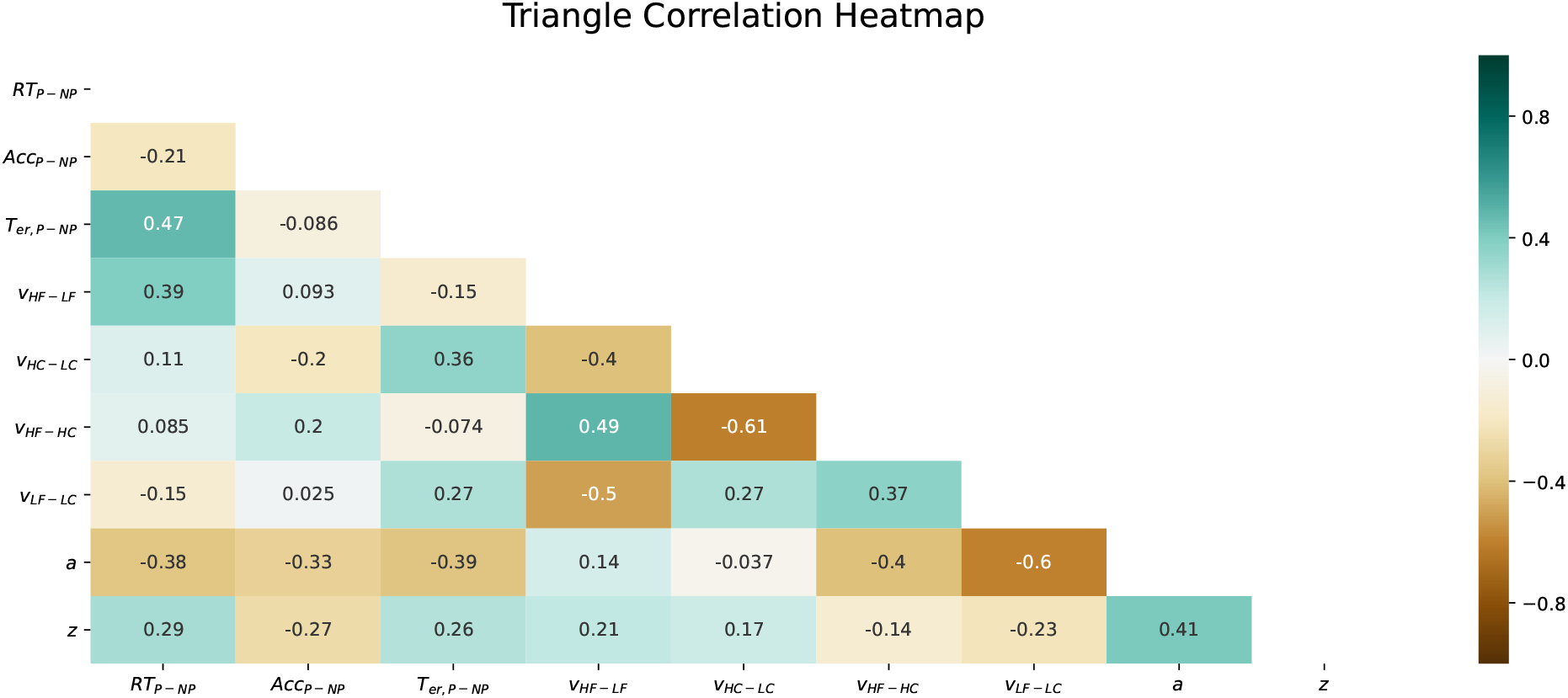
The heatmap matrix correlation of the group-level parameters across conditions. *RT*_*P−NP*_: RT of prioritization minus RT of Non-prioritization, *Acc*_*P−NP*_: Accuracy of prioritization minus Accuracy of Non-prioritization, *T*_*er,P−Np*_: *T*_*er,P*_ minus *T*_*er,NP*_,, *V*_*HF−LF*_ : *V*_*HF*_ minus *V*_*LF*_, *V*_*HC−LC*_: *V*_*HC*_ minus *V*_*LC*_, *V*_*HF−HC*_: *V*_*HF*_ minus *V*_*HC*_, *V*_*LF−LC*_: *V*_*LF*_ minus *V*_*LC*_

### 2.3 Central N2 subcomponent and alpha lateralization

The ERPs calculated from central electrodes C1/2/3/4 are shown in Figure 5. In this figure, N2nc and N2ni are highlighted as gray colors in Figure 5a and Figure 5b respectively. These gray highlights are based on the significant t-test with FDR correction described previously. Figure 5a shows the grand-average across participants of contralateral and neutral ERPs and the difference between them, time-locked to cue onset. FDR correction over samples of the time window found two distinct N2nc subcomponents at central electrode sites with (the mean p = 0.018 < 0.05 for 190-235 msec and the mean p = 0.024 < 0.05 for time windows 260-315 msec after stimulus onset). Figure 5b presents the grand-average of signal ipsilateral and neutral and also ipsilateral minus neutral. As a result, the FDR correction over samples of the time window revealed an N2ni subcomponent at the central electrode sites with (mean p = 0.022 < 0.05 for time window 210-270 msec after stimulus onset).

**Figure 5.**
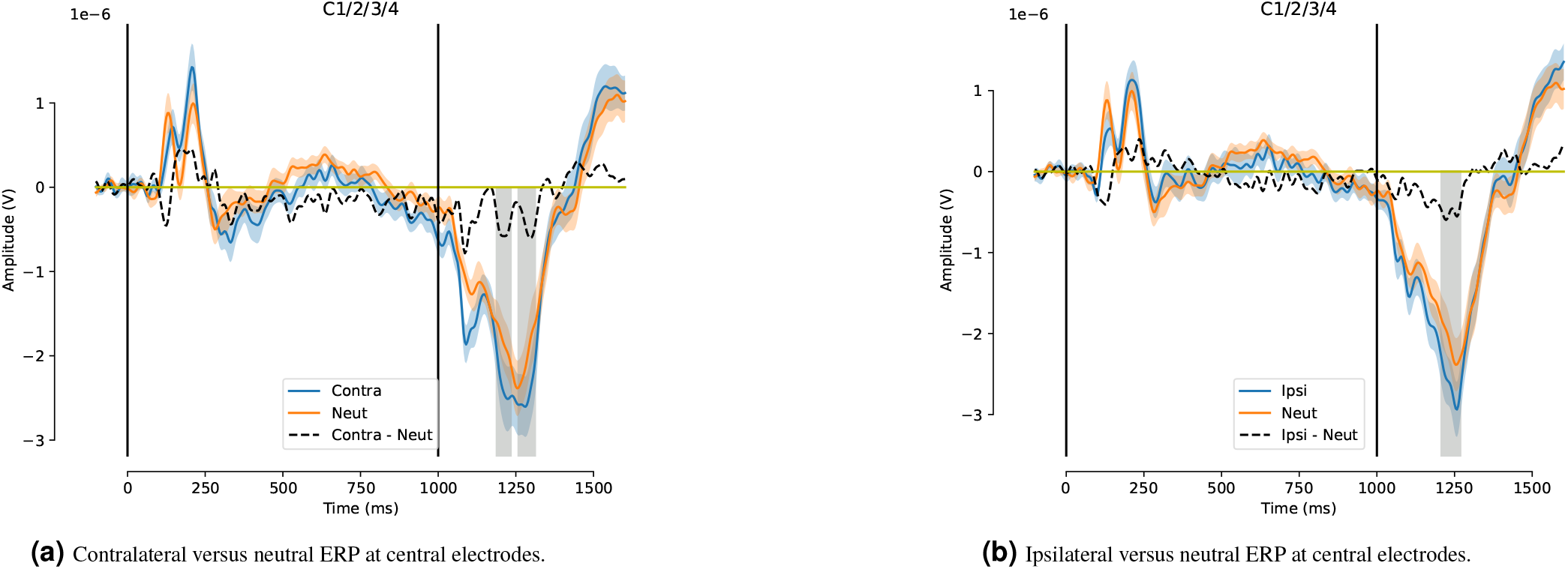
N2nc and N2ni subcomponents at central electrodes C1/2/3/4 highlighted by the shaded gray bars. (a) The grand-average across participants of contralateral and neutral ERPs as well as the N2nc (contralateral minus neutral) are displayed. The significant windows in gray are based on t-tests with *α* = 0.05 and a FDR correction within a time window from 150 msec to 500 msec after stimulus onset are illustrated by the gray color. Specifically the gray color indicates the mean p < 0.05 for 190-235 msec and the mean p < 0.05 for 260-315 msec windows after stimulus onset (i.e. 1190-1235 msec 1260-1315 after cue onset). (b) The grand-average ipsilateral and neutral ERPs as well as the N2ni (ipsilateral minus neutral) are displayed. In addition, the significant windows with on FDR correction are illustrated by the gray color. Specifically the gray color indicates the mean p < 0.05 for 210-270 msec time window after stimulus onset (i.e. 1210-1270 after cue onset). In each figure, the first and second vertical lines show the time point of the spatial cue (single-sided or double-sided arrows) and the stimulus presence (face or house) respectively. The blue and orange shaded regions identify the standard error of each ERP.

Figure 6 presents a grand-average of the single-trial time–frequency representation asymmetric of alpha power from 8 Hz to 13 Hz at posterior-occipital sites (PO3/4/7/8). At first, we considered the ±50 msec around of 50% FAL (237-337 msec after stimulus onset) between -100 msec to 600 msec after stimulus in the Anc signal. Then, by applying a one-sample t-test in the mean portion of Anc, we found a significant effect with (t = 2.59, p = 0.022 < 0.05), as shown in Figure 6a with the gray color. Also, we examined the same analysis in the Ani signal and a time window (155-255 msec after stimulus onset) was found to be significant. Therefore, using a one-sample t-test in the mean portion of Ani, we had a significant effect with (t = 2.27, p = 0.040 < 0.05) which are shown by the gray color, see Figure 6b.

**Figure 6.**
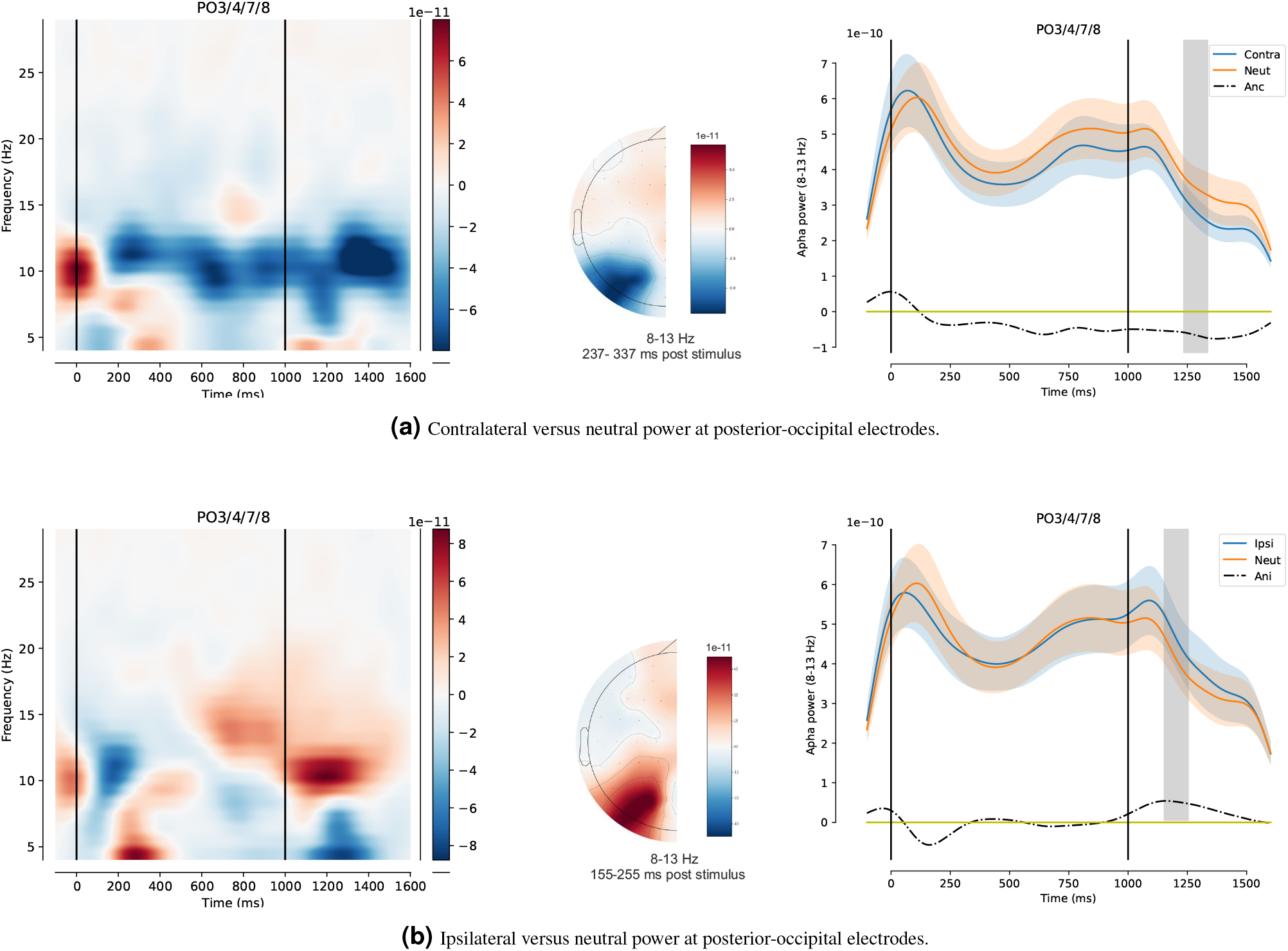
Time–frequency lateralization. (a) The grand-average of contralateral minus neutral single-trial time–frequency wavelets are indicated at electrodes PO3/4/7/8 on the left hand side. In addition, the mean of contralateral and neutral alpha power from 8 Hz to 13 Hz as well as the outcome of (contralateral minus neutral) are displayed on the right hand. The ±50 msec around of 50% FAL on the time window from -100 msec to 600 msec of the stimulus onset was achieved in the 237-337 msec window after the stimulus onset. The topographic map is based on differences of contralateral minus neutral alpha oscillation in the significant time window. (b) The grand-average of ipsilateral minus neutral single-trial time–frequency wavelets are also shown at the posterior-occipital electrodes on the left hand side. Moreover, the mean of contralateral and neutral alpha power as well as the outcome of (ipsilateral minus neutral) are displayed on the right hand side. Therefore, the result of FAL was achieved 155-255 msec after the stimulus onset. The topographic map is based on differences of ipsilateral minus neutral alpha oscillation in the significant time window. The first and second vertical lines show in the right and left plots indicate cue and stimulus occurrences respectively, and shaded regions identify the standard error.

### 2.4 Regression analysis

Tables 1 and 2 show the result of model fitting for both of contralateral versus neutral component and ipsilateral versus neutral component as the difference between informative versus uninformative neutral activity. Mean response times were predicted by both the N2nc with time window 260-315 msec post-stimulus and Anc with the time window 237-337 msec post-stimulus (*R*^2^=0.679,F(2,13)=11.66, p = 0.00192 <0.01). There was no significant effect for N2nc with time window 190-235 post-stimulus, so we disregarded it from further analysis. In this regression model, the main effect of Anc was highly significant (t=3.377, p=0.006<0.01) and the main effect of N2nc was not significant (t=1.549, p=0.150). However the Pearson correlation coefficient disclosed the relationship between mean RT and N2nc (r=0.589, p = 0.026 < 0.05).

**Table 1.** Separate multiple linear regressions between the dependent variables (non-decision time of prioritization minus non-prioritization, RT and accuracy) and predictors (N2nc and Anc), FDR correction for two different N2nc, t-test statistics for each Beta parameters, confidence intervals, and standard errors.

| Output | $T_{er,P-NP}$ | | $RT_{P-NP}$ | | $Acc_{P-NP}$ | |
| --- | --- | --- | --- | --- | --- | --- |
| Predictors | $\beta$ (SE) [95% CI] | t | $\beta$ (SE) [95% CI] | t | $\beta$ (SE) [95% CI] | t |
| Intercept | -0.0046(0.009)<br>[-0.024 0.015] | -0.523<br>p=0.612 | -3.5859(6.836)<br>[-18.63 11.46] | t=-0.525<br>p=0.610 | 0.1295(1.249)<br>[-2.621 2.880] | t=0.104<br>p=0.919 |
| N2nc | 4.712e+04*(1.72e+04)<br>[9354.840 8.49e+04]<br>$BF_{10} = 4.361$ | t=2.746<br>p=0.019 | 2.048e+07(1.32e+07)<br>[8.62e+06 4.96e+07]<br>$BF_{10} = 2.674$ | t= 1.549<br>p=0.150 | -2.999e+06(2.42e+06)<br>[-8.32e+06 2.32e+06]<br>$BF_{10} = 0.684$ | t=-1.241<br>p=0.241 |
| Anc | -3.485e+07(6.92e+07)<br>[-1.87e+08 1.17e+08]<br>$BF_{10} = 0.516$ | t=-0.504<br>p=0.624 | 1.8e+11**(5.33e+10)<br>[6.27e+10 2.97e+11]<br>$BF_{10} = 29.217$ | t=3.377<br>p=0.006 | 5.19e+09(9.74e+09)<br>[-1.63e+10 2.66e+10]<br>$BF_{10} = 0.446$ | t=0.533<br>p=0.605 |
| $R^2$ | 0.428 | | 0.679 | | 0.123 | |
| F-statistic | F(2,13) = 4.114, p = 0.0464<br>$BF_{10} = 2.036$ | | F(2,13) = 11.66, p=0.00192<br>$BF_{10} = 21.83$ | | F(2,13) = 0.7701, p = 0.486<br>$BF_{10} = 0.415$ | |

$T_{P-NP} = T_{er}(\text{yes}) - T_{er}(\text{no})$ , $RT_{P-NP} = RT(P) - RT(NP)$ , $Acc_{P-NP} = \text{accuracy}(P) - \text{accuracy}(NP)$ , SE = standard error, CI = confidence interval. $R^2$ and regression coefficients are given for separate linear regression models, yes= prioritization, no= no prioritization. Asterisks indicate significant p value as \*p<0.05, \*\*p<0.01.

**Table 2.** Separate multiple linear regressions between the dependent variables (non-decision time of prioritization minus non-prioritization, RT and accuracy) and predictors (N2ni and Ani), FDR correction for two different N2nc, t-test statistics for each Beta parameters, confidence intervals, and standard errors.

| Output | $T_{er,P-NP}$ | | $RT_{P-NP}$ | | $Acc_{P-NP}$ | |
| --- | --- | --- | --- | --- | --- | --- |
| Predictors | $\beta$ (SE) [95% CI] | t | $\beta$ (SE) [95% CI] | t | $\beta$ (SE) [95% CI] | t |
| Intercept | -0.0194(0.013)<br>[-0.047 0.008] | t=-1.548<br>p=0.150 | -11.6572(10.140)<br>[-33.976 10.662] | t=1.1700(1.326)<br>p=0.275 | t=0.882<br>[-1.749 4.089] | p=0.397 |
| N2ni | 8286.7188(2.52e+04)<br>[-4.71e+04 6.36e+04]<br>$BF_{10} = 0.466$ | t=0.329<br>p=0.748 | 8.253e+06(2.04e+07)<br>[-3.66e+07 5.32e+07]<br>$BF_{10} = 1.333$ | t= 0.405<br>p=0.694 | 1.82e+06(2.67e+06)<br>[-4.05e+06 7.69e+06]<br>$BF_{10} = 0.456$ | t=0.682<br>p=0.509 |
| Ani | 1.225e+07(1.22e+08)<br>[-2.56e+08 2.81e+08]<br>$BF_{10} = 0.450$ | t=0.100<br>p=0.922 | -1.643e+11(9.89e+10)<br>[-3.82e+11 5.35e+10]<br>$BF_{10} = 3.312$ | t=-1.661<br>p=0.125 | 1.703e+10(1.29e+10)<br>[-1.14e+10 4.55e+10]<br>$BF_{10} = 0.698$ | t=1.316<br>p=0.215 |
| $R^2$ | 0.012 | | 0.387 | | 0.141 | |
| F-statistic | F(2,13) = 0.0674, p =0.94<br>$BF_{10} = 0.272$ | | F(2,13) = 3.469, p =0.0679<br>$BF_{10} = 1.560$ | | F(2,13) = 0.7701, p =0.434<br>$BF_{10} = 0.447$ | |

$T_{P-NP} = T_{er}(P) - T_{er}(NP)$ , $RT_{P-NP} = RT(P) - RT(NP)$ , $Acc_{P-NP} = accuracy(P) - accuracy(NP)$ , SE = standard error, CI = confidence interval. $R^2$ and regression coefficients are given for separate linear regression models, yes= prioritization, no= no prioritization. Asterisks indicate significant p value as \*p<0.05, \*\*p<0.01.

Non-decision time differences explained by N2nc and Anc (*R*^2^=0.428, F(2,13)=4.114, p = 0.0464< 0.05). In addition, the main effect of N2nc was significant (t=2.746, p = 0.019 <0.05), and the main effect of Anc was not significant (t=-0.504, p=0.624). However accuracy was not predicted by N2nc and Anc (*R*^2^=0.123, F(2,13)=0.7701, p=0.486).

For regressions with ipsilateral versus neutral regressors, there were also no significant models, see Table 2. The visualizations of the results of these regressions are reported by Component-Component plus Residual (CCPR) for the non-decision time parameter and RT regression models with N2nc and Anc, and non-decision time regression model with N2ni and Ani, see Figure 7.

**Figure 7.**
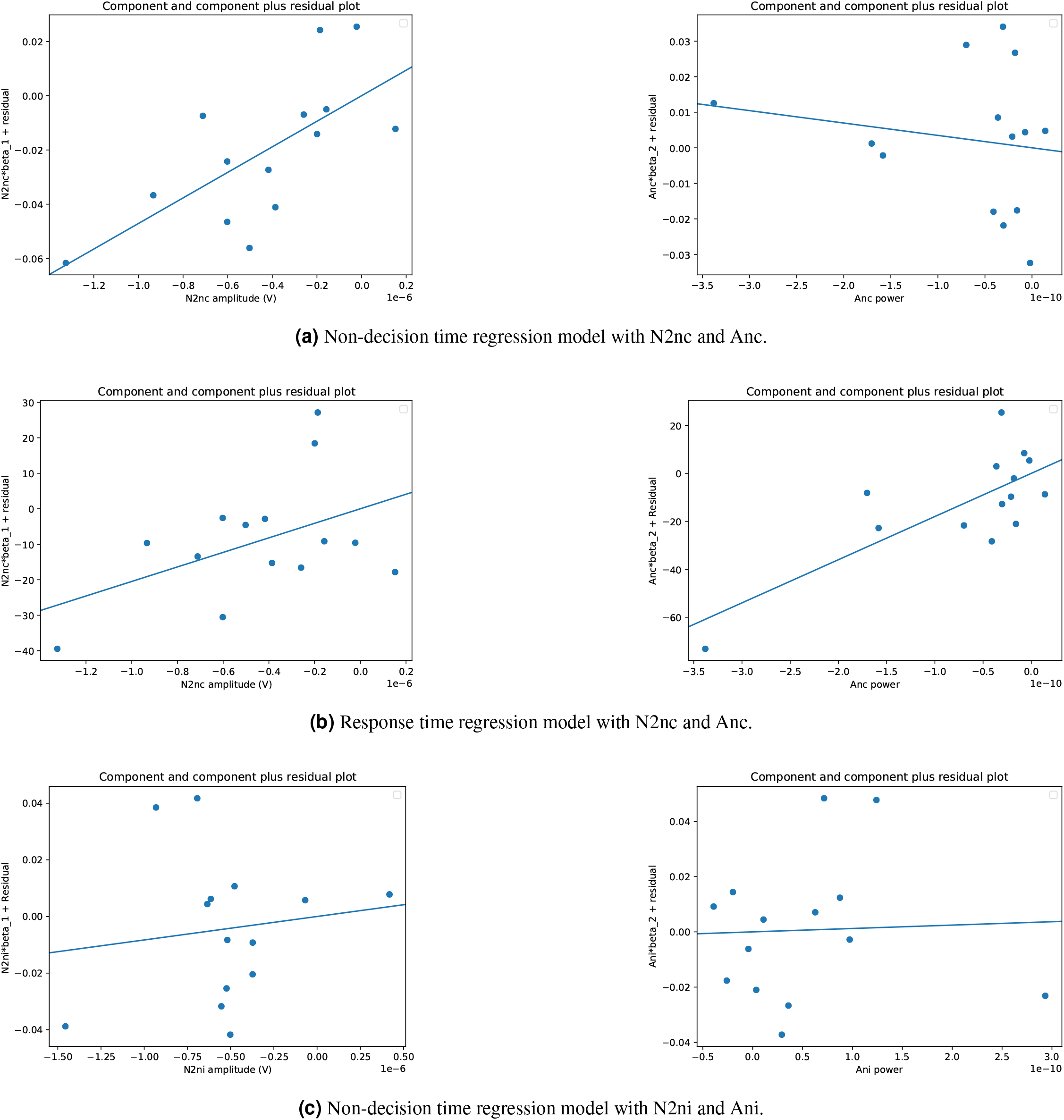
Component-Component plus Residual plots, a. For the non-decision parameter model, the main effect of the N2nc amplitude is significant and Anc was no significan, b. For the RT time regression model, the main effect of the Anc power is significant and the N2nc was not significant. But, the difference RT was also highly correlated with N2nc by Pearson correlation coefficient which had not seen in the regression model, c. The N2i (t=0.329, p=0.748) and Ani were not significant effects in non-decision time. Moreover, all regressors were not significant effects in accuracy mode in both RT and non-decision time regression.

## 3 Discussion

This work investigated how spatial prioritization cues lead to faster response times than non-spatial prioritization cues in the perceptual decision making tasks. In order to explain the elaborate framework of the observed behavior, cognitive process, and observed electrophysiology during top-down intention, we focused on the neural contribution of post-stimulus N2 subcomponents and alpha power. Contralateral minus uninformative N2 subcomponents and alpha power predicted the behavioral data and relevant DDM parameters. We differentiated between five competing models: 1. Spatial attention may not shift resources related to decision-making. 2. Spatial attention might affect the information accumulation process. 3. Spatial attention could be related to the speed of decoding or motor response (non-decision time). 4. Spatial attention could lead to a bias to the presented stimulus. 5. Spatial attention could shift the amount of information that is needed to decide. We found Model 3 (model_*t*_) best described the data as measured by DIC and *R*^2^. In addition, we tested whether lateralized central sites relating to motor response and parieto-occipital relating to decoding stimulus reflected individual differences in model parameters. Specifically, we investigated the contribution of alpha power lateralization and N2 sub-component amplitudes to individual differences in spatial attention in perceptual decision making. Our study revealed that both contralateral N2 and alpha power regressors could predict the non-decision parameter of the model_*t*_ and RT associated with the spatial cue, see Table 1. Whereas, ipsilateral biomarkers could not predict non-decision parameter of the model_*t*_ and RT associated with the spatial cue, see Table 2.

We found that only the difference between contralateral and neutral components which could predict the difference in non-decision times (T_*er,P−NP*_ = T_*er*_(P) - T_*er*_(NP)) and RT (RT_*P−NP*_ = RT(P) - RT(NP)) between prioritization and non-prioritization. This finding confirms that the mechanism of contralateral oscillations to the attended spatial location is more important than ipsilateral oscillation. Most previous works (^5, 25, 26, 29^) have concentrated on the difference of contralateral and ipsilateral deflections and have not explored which signals, by themselves, could predict relevant the DDM parameters associated with spatial attention. In the current study we also found a significant relationship between non-decision time and contralateral minus neutral activity (^5, 22^).

### 3.1 N2nc and Anc band predict response time

Behavioral data revealed a significant effect of spatial prioritization relating to non-prioritization. Therefore, in order to decipher the mechanism of the difference, a multiple regression model using N2nc (the time measurement 260-315 msec) and Anc band as regressors was fit. This model could explain the difference RT between prioritization and non-prioritization, see Table 1. The effect of N2nc and Anc coefficients were displayed in Figure 7b. Also, the components of N2ni and Ani together could not explain the variance of the difference RT, see Table 2. This result shows that contralateral activity could encode the different spatial cues in comparison with ipsilateral activity. According to post-stimulus of figure 5, both N2 subcomponent contralateral and ipsilateral ERP were more negative than neutral ERP in line with latter findings (^22, 26^).

Post-stimulus contralateral alpha power was more negative than neutral alpha power, but ipsilateral alpha power was more positive than neutral alpha power, see 6. Negative Anc suggests that the alpha power contralateral to the attend visual field decreased because participants’ brains were made more sensitive to decode the stimulus from the attended area. On other hand, positive Ani shows that alpha power ipsilateral to the attend visual field increased because it is thought the participants’ brains to sought to attenuate the non-attended spatial location (^17, 78^). In our research we separated these two mechanisms in order to understand the role of spatial attention encoded by EEG on perceptual decision making. The regression and statistical analyses proved that the role Anc and N2nc are more significant to spatial attention. That is N2nc and Anc led to the more differences in RT associated with spatial attention.

### 3.2 N2nc amplitude predicts non-decision times

We also explored the role of alpha lateralization and N2nc in visual perceptual decision making assuming a cognitive model, a DDM. We found that the difference in RT of prioritization (informative cue conditions) and non-prioritization (uninformative cue conditions) originated from the difference of non-decision time parameter based on the nested model comparisons. We show that N2nc and Anc predicted the non-decision parameter based on the nested model comparison, see Table 1. However the main effect of N2nc was significant, but Anc was not significant, see Figure 7a. N2ni and Ani could not predict non-decision time parameter, see Table 2 and Figure 7c. Thus the N2nc might reflect the non-decision parameter that influences RT, while the Anc might reflect RTs without a clear cognitive parameter correlate.

The role of N2nc was easier to interpret as mediating the non-decision parameter difference. It also shows that the contralateral activity was superior to elucidate the effect of the spatial cue in comparison with ipsilateral activity. Therefore, the larger differences in non-decision time measured by N2nc, the larger the differences of non-decision time relating to spatial attention.

## 4 Conclusion and future works

The non-decision time parameter reflects the spatial cue in perceptual decision making based on our findings. We also found that the amplitude of N2nc at the central site predicted RT as well as the non-decision time due to spatial prioritization across participants. The Anc power oscillations at parieto-occipital electrodes only predicted RT of spatial prioritization across participants and did not predict non-decision time across participants. In addition, neither the N2ni amplitudes and Ani power could predict RT and non-decision time differences between spatial prioritization and non-prioritization across participants. Therefore, the role of contralateral activity is shown to allocate the brain’s resources in order to aid the decision making process, in comparison to ipsilateral activity. Larger N2nc led to larger differences in non-decision time associated with spatial attention, while both larger N2nc or Anc differences led larger differences of mean RTs associated with spatial attention. In addition, the non-decision time parameter might explain the modulation of the attentional orienting during decision making process.

For future work, different joint-modeling techniques such as the Integrative, Directed, and Covariance approaches should be used in order to provide a moment-by-moment descriptions of the neural data and a trial-by-trial descriptions of behavioral data. Moreover, we combining latent processes and the dynamics underlyings of varied modalities (e.g. bahavioral data, EEG, fMRI and so forth) could yield more insight into the neurocognitive role of spatial attention (^28, 79–83^).

## Acknowledgement

All data used in this research is publicly available in the open science framework (https://osf.io/q4t8k/). The authors are very grateful to Dirk Ostwald and his colleagues that made this data available. MDN was supported in part by US NSF grants (1658303 and 1850849).

## Supplementary Material

### Model fitting and Model comparison

In this paper, five possible models were run to give us appropriate intuition of spatial prioritization. To compare which model is better to fit observed data, our analysis was equipped with two powerful weapons called DIC score (lower is better) and *R*^2^ (near to one is better). It should be noted that a difference in DIC scores of 15 and above is considered meaningful^70, 75^. Therefore, the result of model comparison analysis reveals that model_*t*_ provided the best model fit so that it received the smallest DIC and largest *R*^2^, so non-decision time parameter (*T*_*er*_) interpret the manipulation of spatial prioritization, see Table 3.

**Table 3.**
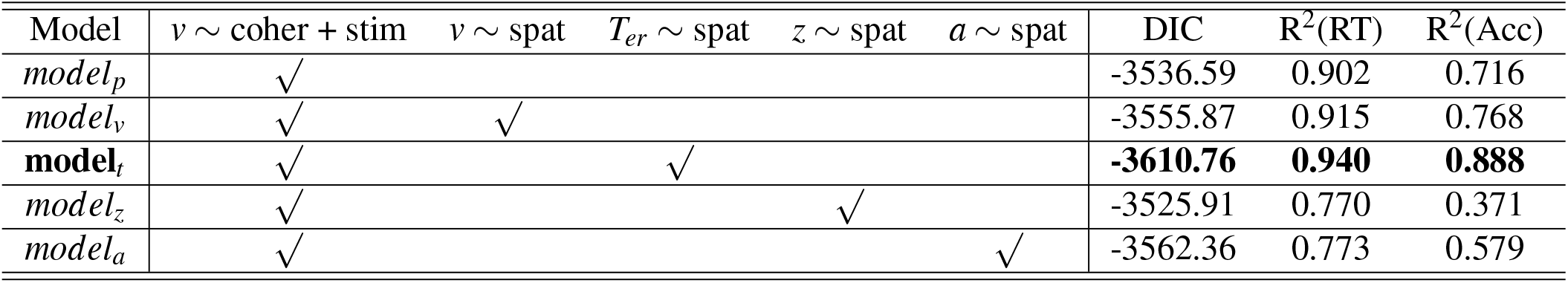
Model comparison based on DIC for all model variants tested in this study and R2 metrics for RT and accuracy in the group level. coher, coherence; stim, stimulus; spat, spatial attention; RT, response time; Acc, accuracy.

To assess model convergence (i.e. model_*t*_), the MCMC sampling traces of the group posteriors distribution for these parameters which show converged chain procedure during parameters estimation were utilized, see Figures 8, 9 and 10. There were horizontal lines (convergence) in the trace and no jump in the auto-correlation plots, and the group mean posteriors were smooth with no jagged signs.

**Figure 8.**
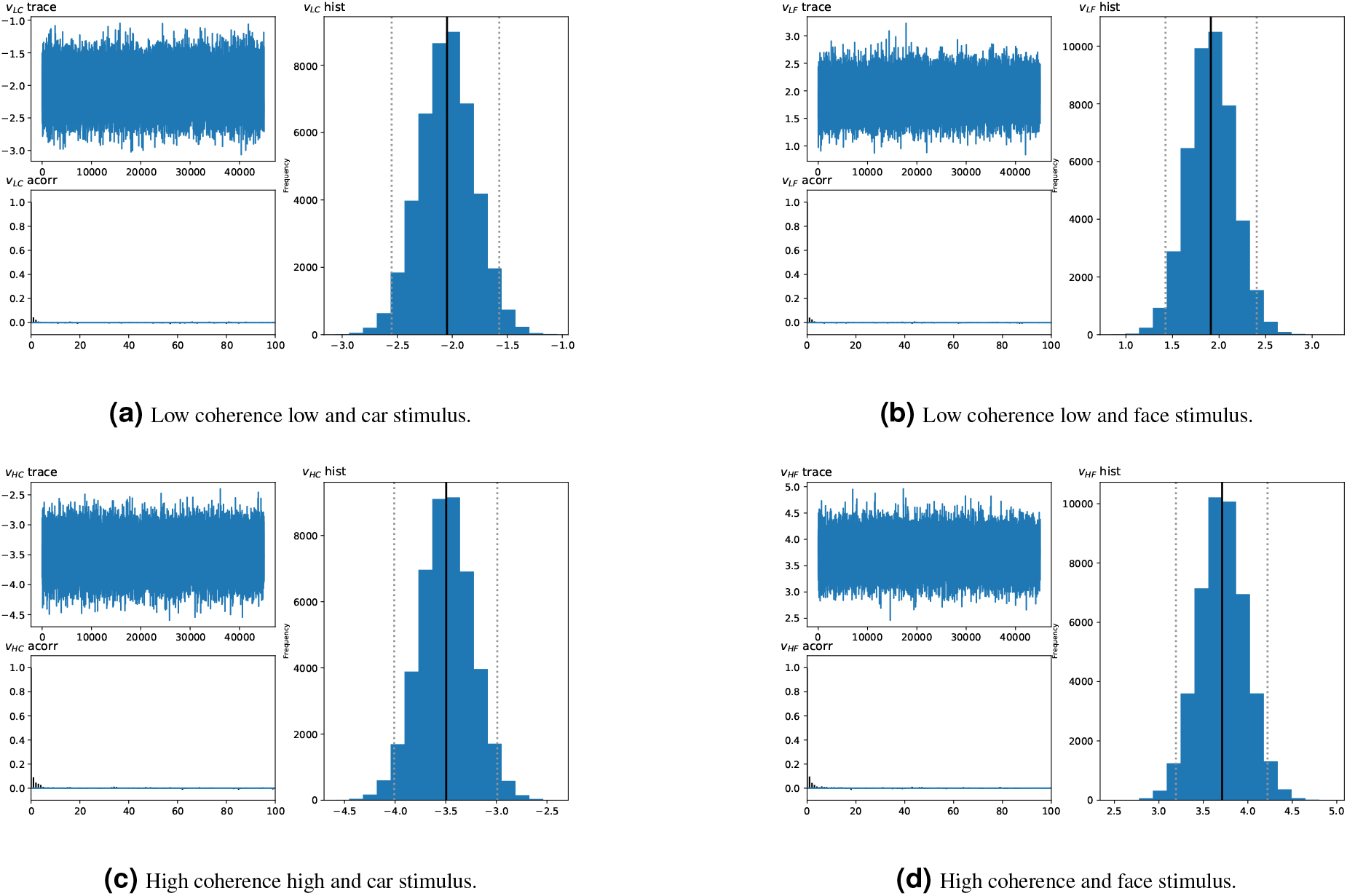
Plots of *model*_*t*_ convergence evaluation with the trace, auto-correlation, and histogram posterior distribution of drift rate parameter for the low coherence (upper half) and high coherence (bottom half) conditions of both types of car and face stimuli.

**Figure 9.**
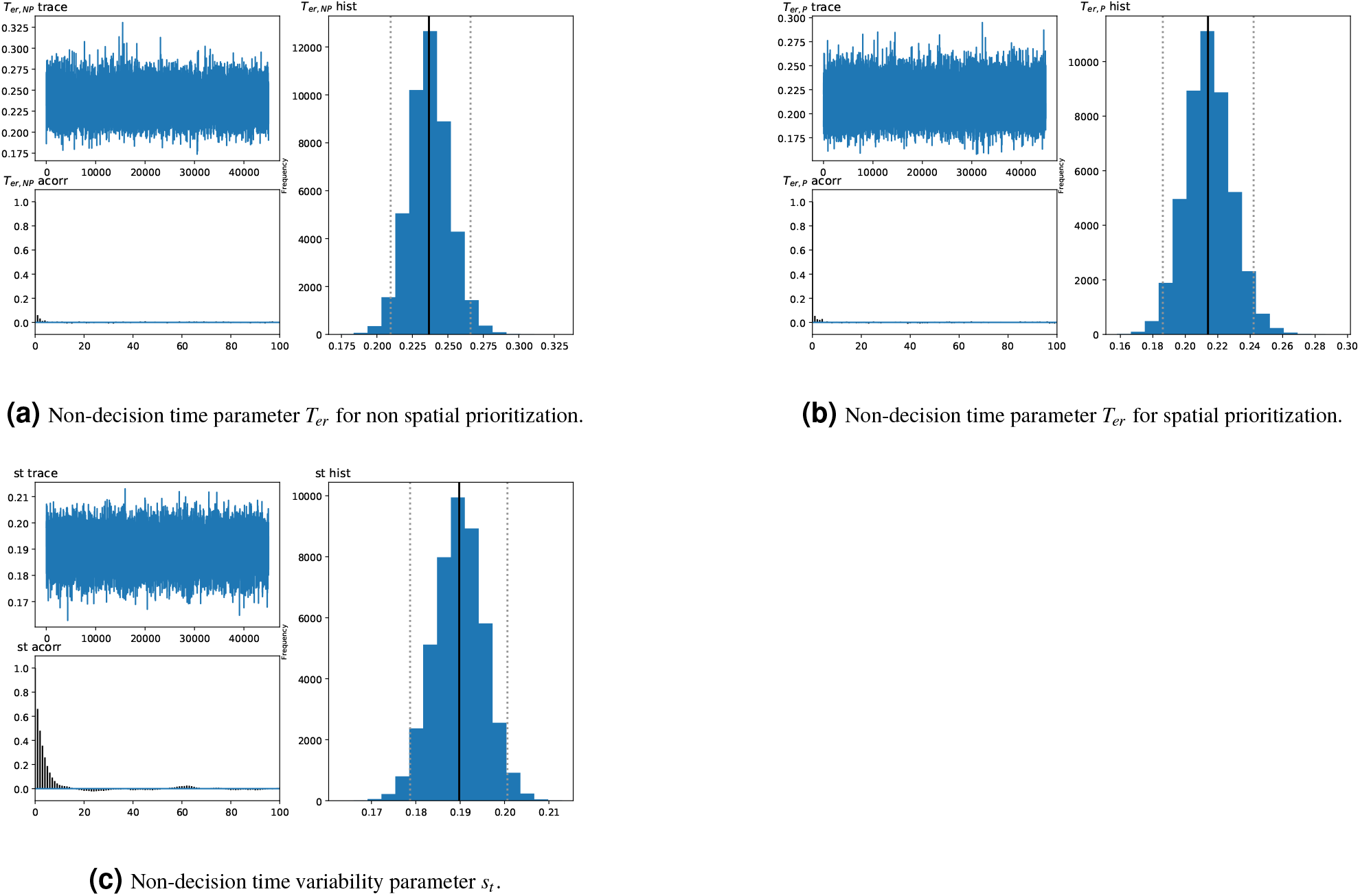
Plots of *model*_*t*_ convergence evaluation with the trace, auto-correlation, and histogram posterior distribution of non-decision time parameter for the prioritization (top left) and no prioritization coherence (top right) conditions. Non-decision time variability is also reported in the bottom half.

**Figure 10.**
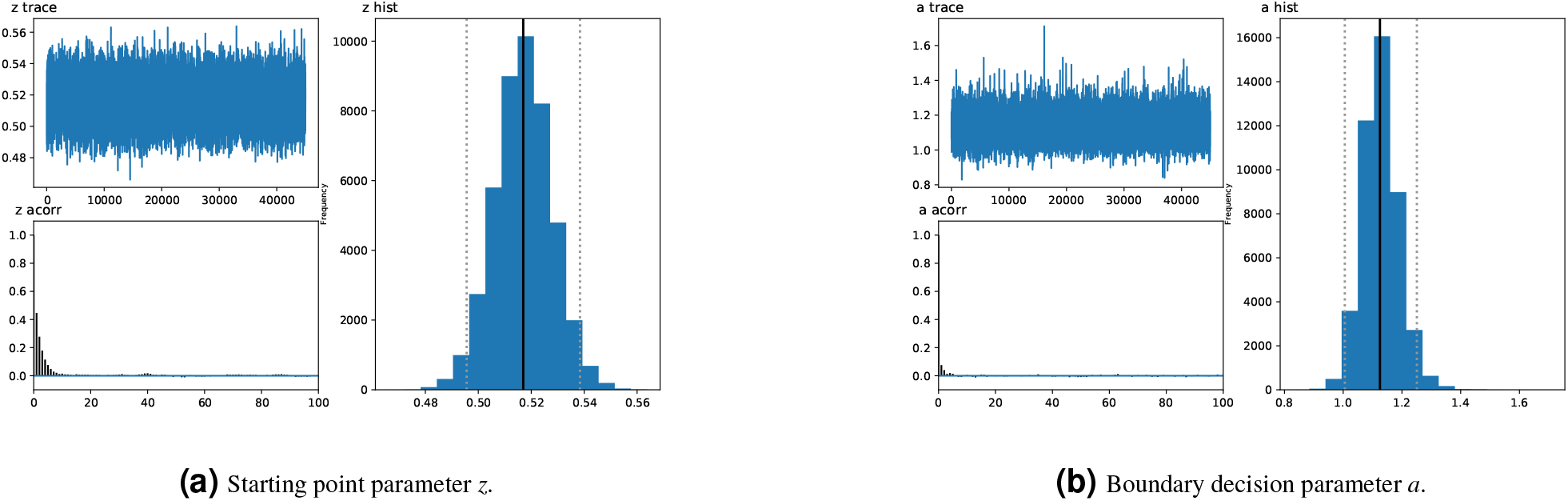
Plots of *model*_*t*_ convergence evaluation with the trace, auto-correlation, and histogram posterior distribution of (a) starting point parameter and (b) boundary decision parameter.

*R-hat values for all models*: The convergence of the Markov chains was assessed by calculating the R-hat Gelman-Rubin statistic (^74^) to ensure that the models had properly converged, which compares between-chain and within-chain variance. The model of “model_*t*_” showed superior convergence and the same stationary distribution with four chains, based on R-hat values under 1.0001 as follow:

**model**_*t*_:

’*T*_*er,NP*_’:0.9999935100922528,

’*T*_*er,P*_’: 0.9999916748628642,

’*v*_*HC*_’: 1.0000426841659067,

’*v*_*HF*_ ‘: 0.999991945644348,

’*v*_*LC*_’: 1.0000080181533444,

’*v*_*LF*_ ‘: 1.0000028712137732,

’*a*’: 0.9999968891198591,

’*st*’: 1.0000144779358509,

’*z*’: 1.0000630870848732,

#### Plot of posterior predictive against observed data

To assess the model fit, we compared posterior model predictions with the observed data. In fact, posterior predictive checks of the best model were conducted by averaging 500 simulations generated from the model’s posterior to confirm it could reliably reproduce patterns in the observed data. There was a good agreement between the observed data and the model predictions across conditions in all conditions. Overall, the model is able to capture the overall shape of response times across the four different conditions, see Figure 11.

**Figure 11.**
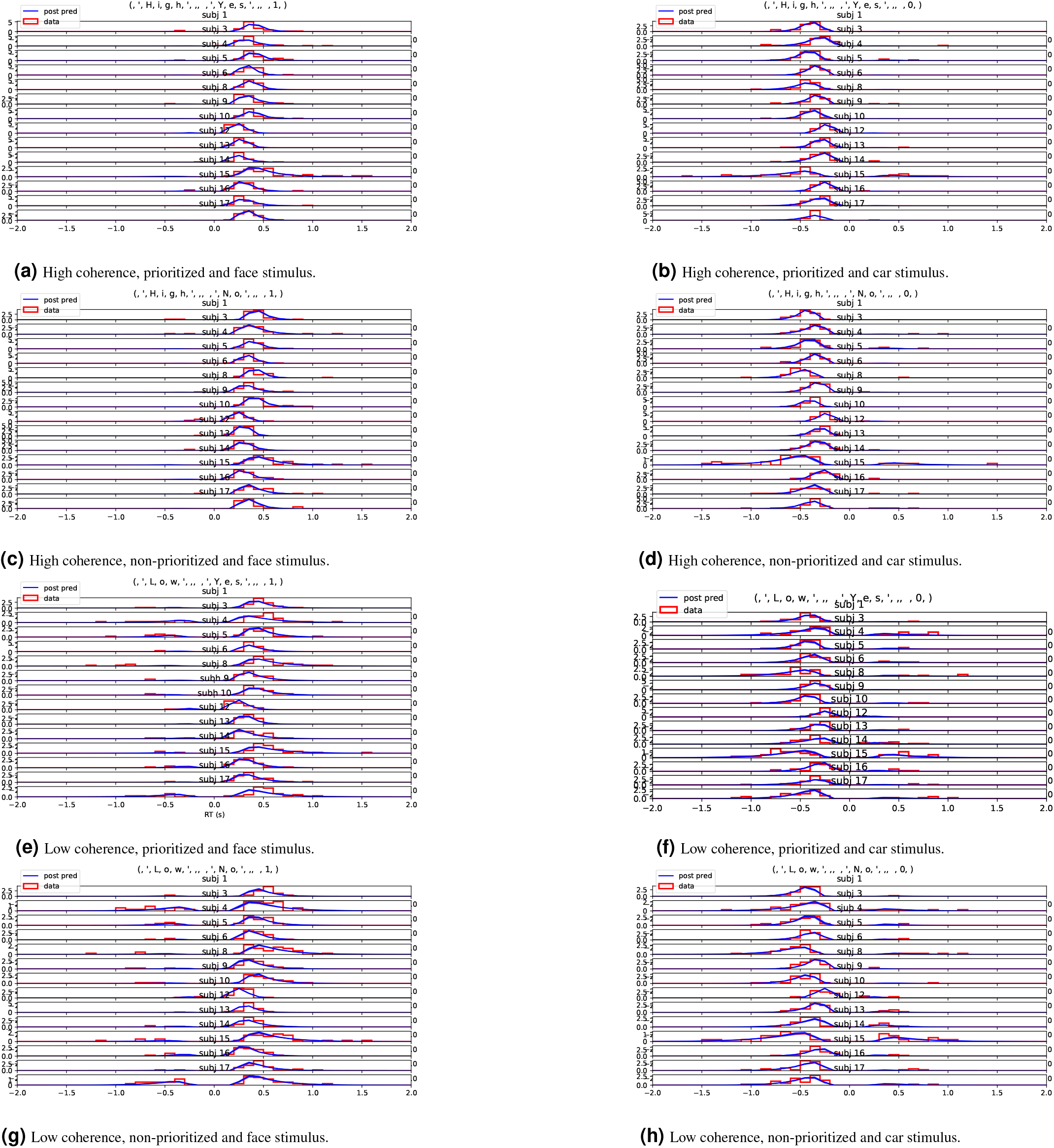
The histogram of *model*_*t*_ shows observed response time distribution for all participants across all conditions (positive vs. negative results on the x-axis correspond to face vs. car stimuli). The line plots the results from the posterior predictive simulation.

## References

1. Gold, J. I. & Shadlen, M. N. The neural basis of decision-making. Annu. Rev. Neurosci. 30, 535–574 (2007).

2. Gherman, S. & Philiastides, M. G. Human vmpfc encodes early signatures of confidence in perceptual decisions. eLife 7, 1–18 (2018).

3. Shadlen, M. N. & Kiani, R. Decision making as a window on cognition. Nature 80, 791–806 (2013).

4. Philiastides, M. G., Ratcliff, R. & Sajda, P. Neural representation of task difficulty and decision making during perceptual categorization: a timing diagram. J. Neurosci. 26, 8965–8975 (2006).

5. Klatt, L. I. et al.. Unraveling the relation between EEG correlates of attentional orienting and sound localization performance: a diffusion model approach. J. Cogn. Neurosci. 32, 945–962 (2020).

6. Krajbich, I., Lu, D., Camerer, C. & Rangel, A. The attentional drift-diffusion model extends to simple purchasing decisions. Front. Psychol. 3, 193 (2012).

7. Nunez, M. D., Vandekerckhove, J. & Srinivasan, R. How attention influences perceptual decision making: Single-trial eeg correlates of drift-diffusion model parameters. J. mathematical psychology 76, 117–130 (2017).

8. S. Gluth and M. S. Spektor and J. Rieskamp. Value-based attentional capture affects multi-alternative decision making. eLife 7, e39659 (2018).

9. Nunez, M. D., Srinivasan, R. & Vandekerckhove, J. Individual differences in attention influence perceptual decision making. Front. psychology 6, 18 (2015).

10. Ostwald, D., Porcaro, C., Mayhew, S. D. & Bagshaw, A. P. EEG-fMRI based information theoretic characterization of the human perceptual decision system. PLoS ONE 7, e33896 (2012).

11. Yeshurun, Y. & Carrasco, M. Spatial attention improves performance in spatial resolution tasks. Vis. Res. 39, 293–306 (1999).

12. Posner, M. I. Orienting of attention. The Q. journal experimental psychology 32, 3–25 (1980).

13. Sagar, V., Sengupta, R., Devarajan & Sridharan. Dissociable sensitivity and bias mechanisms mediate behavioral effects of exogenous attention. Sci. Rep. 9, 1–13 (2019).

14. Imani, E., Harati, A., Pourreza, H. & Goudarzi, M. M. Brain-behavior relationships in the perceptual decision-making process through cognitive processing stages. Neuropsychologia 155, 107821 (2021).

15. Foster, J. J., Sutterer, D. W., Serences, J. T., Vogel, E. K. & Awh, E. Alpha-band oscillations enable spatially and temporally resolved tracking of covert spatial attention. Psychol. Sci. 28, 929–941 (2017).

16. Li, Y., Lou, B., Gao, X. & Sajda, P. Post-stimulus endogenous and exogenous oscillations are differentially modulated by task difficulty. Front. Hum. Neurosci. 7, 1–10 (2013).

17. Rihs, T. A., Michel, C. M. & Thut, G. Mechanisms of selective inhibition in visual spatial attention are indexed by α-band eeg synchronization. Eur. J. Neurosci. 25, 603–610 (2007).

18. van Schouwenburg, M. R., Zanto, T. P. & Gazzaley, A. Spatial attention and the effects of frontoparietal alpha band stimulation. Front. human neuroscience 10, 658 (2017).

19. Praamstra, P., Boutsen, L. & Humphreys, G. W. Frontoparietal control of spatial attention and motor intention in human eeg. J. neurophysiology 94, 764–774 (2005).

20. Haegens, S., Luther, L. & Jensen, O. Somatosensory anticipatory alpha activity increases to suppress distracting input. J. cognitive neuroscience 24, 677–685 (2012).

21. Bernier, P. M., Whittingstall, K. & Grafton, S. T. Differential recruitment of parietal cortex during spatial and non-spatial reach planning. Front. human neuroscience 11, 249 (2017).

22. Praamstra, P. Prior information of stimulus location: effects on erp measures of visual selection and response selection. Brain research 1072, 153–160 (2006).

23. Amenedo, E., Lorenzo-Lopez, L. & Pazo-alvarez, P. Response processing during visual search in normal aging: the need for more time to prevent cross talk between spatial attention and manual response selection. Biol. psychology 91, 201–211 (2012).

24. Praamstra, P. & Oostenveld, R. Attention and movement-related motor cortex activation: a high-density eeg study of spatial stimulus–response compatibility. Cogn. Brain Res. 16, 309–322 (2003).

25. Cespon, J., Galdo-Alvarez, S. & Diaz, F. Cognitive control activity is modulated by the magnitude of interference and pre-activation of monitoring mechanisms. Sci. reports 6, 1–11 (2016).

26. Gamble, M. L. & Luck, S. J. N2ac: An erp component associated with the focusing of attention within an auditory scene. Psychophysiology 48, 1057–1068 (2011).

27. Loughnane, G. M. et al.. Target selection signals influence perceptual decisions by modulating the onset and rate of evidence accumulation. Curr. Biol. 26, 496–502 (2016).

28. Nunez, M. D., Gosai, A., Vandekerckhove, J. & Srinivasan, R. The latency of a visual evoked potential tracks the onset of decision making. Neuroimage 197, 93–108 (2019).

29. Luck, S. J. An introduction to the event-related potential technique. J. Chem. Inf. Model. 53 (2014).

30. Ratcliff, R. & McKoon, G. The diffusion decision model: theory and data for two-choice decision tasks. Neural computation 20, 873–922 (2004).

31. Olianezhad, F., Zabbah, S., Tohidi-Moghaddam, M. & Ebrahimpour, R. Residual information of previous decision affects evidence accumulation in current decision. Front. Behav. Neurosci. 13, 1–12 (2019).

32. Birte U. Forstmann and Roger Ratcliff and Eric-Jan Wagenmakers. Sequential sampling models in cognitive neuroscience: Advantages, applications, and extensions. Annu. review psychology 67, 641–666 (2016).

33. Smith, P. L. Diffusion theory of decision making in continuous report. Psychol. Rev. 123, 425–451 (2016).

34. Ratcliff, R. A theory of memory retrieval. Psychol. review 85, 59 (1978).

35. Stone, M. Models for choice-reaction time. Psychometrika 25, 251–260 (1960).

36. Ratcliff, R., Smith, P. L., Brown, S. D. & McKoon, G. Diffusion decision model: Current issues and history. Trends Cogn. Sci. 20, 260–281 (2016).

37. Evans, N. J. & Wagenmakers, E.-J. Evidence accumulation models: Current limitations and future directions. The Quant. Methods for Psychol. (2020).

38. Evans, Nathan J and Hawkins, Guy E and Brown, Scott D. The role of passing time in decision-making. J. experimental psychology: learning, memory, cognition 46, 316 (2020).

39. Evans, N. J. & Brown, S. D. People adopt optimal policies in simple decision-making, after practice and guidance. Psychon. Bull. & Rev. 24, 597–606 (2017).

40. Evans, N. J., Bennett, A. J. & Brown, S. D. Optimal or not; depends on the task. Psychon. bulletin & review 26, 1027–1034 (2018).

41. Drugowitsch, J., Moreno-Bote, R., Churchland, A. K., Shadlen, M. N. & Pouget, A. The cost of accumulating evidence in perceptual decision making. The J. Neurosci. 32, 3612–3628 (2012).

42. Matzke, D., Dolan, C. V., Logan, G. D., Brown, S. D. & Wagenmakers, E.-J. Bayesian parametric estimation of stop-signal reaction time distributions. J. Exp. Psychol. Gen. 142, 1047 (2013).

43. Gomez, P., Ratcliff, R. & Perea, M. A model of the go/no-go task. J. Exp. Psychol. Gen. 136, 389 (2007).

44. Usher, M. & McClelland, J. L. Loss aversion and inhibition in dynamical models of multialternative choice. Psychol. Rev. 111, 757–769 (2004).

45. P. D. Kvam. A geometric framework for modeling dynamic decisions among arbitrarily many alternatives. J. Math. Psychol. 91, 14–37 (2019).

46. Mallahi-Karai, K. & Diederich, A. Decision with multiple alternatives: Geometric models in higher dimensions - the cube model. J. Math. Psychol. 93, 102294 (2019).

47. Pedersen, M. L., Frank, M. J. & Biele, G. The drift diffusion model as the choice rule in reinforcement learning. Psychon. bulletin & review 24, 1234–1251 (2017).

48. Fontanesi, L., Gluth, S., Spektor, M. S. & Rieskamp, J. A reinforcement learning diffusion decision model for value-based decisions. Psychon. bulletin & review 26, 1099–1121 (2019).

49. D. K. Sewell and H. K. Jach and R. J. Boag and C. A. Van Heer. Combining error-driven models of associative learning with evidence accumulation models of decision-making. Psychon. bulletin & review 26, 868–893 (2019).

50. Gluth, S., Kern, N., Kortmann, M. & Vitali, C. L. Value-based attention but not divisive normalization influences decisions with multiple alternatives. Nat. Hum. Behav. 4, 634–645 (2020).

51. Ratcliff, R. Decision making on spatially continuous scales. Psychol. Rev. 125, 888–935 (2018).

52. Choi, W. & Paik, S.-B. Intrinsic timescales of sensory integration for motion perception. Sci. Reports 9, 1–15 (2019).

53. Voss, A., Rothermund, K. & Voss, J. Interpreting the parameters of the diffusion model: An empirical validation. Mem. & cognition 32, 1206–1220 (2004).

54. Wagenmakers, E.-J., Van Der Maas, H. L. & Grasman, R. P. An ez-diffusion model for response time and accuracy. Psychon. bulletin & review 14, 3–22 (2007).

55. Brown, S. & Heathcote, A. A ballistic model of choice response time. Psychol. review 112, 117 (2005).

56. Usher, M. & McClelland, J. L. The time course of perceptual choice: the leaky, competing accumulator model. Psychol. review 108, 550 (2001).

57. Tillman, G., Van Zandtc, T. & Loganb, G. D. Sequential sampling models without random between-trial variability: the racing diffusion model of speeded decision making. Psychon. bulletin & review (2020).

58. Ratcliff, R. & Smith, P. Modeling simple decisions and applications using a diffusion model. Oxf. Univ. Press. (2015).

59. Voss, A., Nagler, M. & Lerche, V. Diffusion models in experimental psychology: A practical introduction. Exp. Psychology 60, 385 (2013).

60. R. Ratcliff and J. N. Rouder. Modeling response times for two-choice decisions. Psychol. Sci. 9, 347–356 (1998).

61. Ratcliff, R. & Tuerlinckx, F. Estimating parameters of the diffusion model: Approaches to dealing with contaminant reaction times and parameter variability. Psychon. bulletin & review 9, 438–481 (2002).

62. Georgie, Y. K., Porcaro, C., Bagshaw, A. P., Mayhew, S. D. & Ostwald, D. A perceptual decision making eeg / fmri data set. bioRxiv (2018).

63. Gramfort, A. et al.. Meg and eeg data analysis with mne-python. Front. Neurosci. 7 (2013).

64. Nunez, M. D., Nunez, P. L. & Srinivasan, R. Electroencephalography (EEG): neurophysics, experimental methods, and signal processing. In Ombao, H., Linquist, M., Thompson, W. & Aston, J. (eds.) Handbook of Neuroimaging Data Analysis, 175–197, DOI: 10.13140/rg.2.2.12706.63687 (Chapman & Hall/CRC, 2016).

65. Wilsch, A., Mercier, M. R., Obleser, J., Schroeder, C. E. & Haegens, S. Spatial attention and temporal expectation exert differential effects on visual and auditory discrimination. J. cognitive neuroscience 32, 1562–1576 (2020).

66. Van Dijk, H., Schoffelen, J.-M., Oostenveld, R. & Jensen, O. Prestimulus oscillatory activity in the alpha band predicts visual discrimination ability. J. Neurosci. 28, 1816–1823 (2008).

67. Vandekerckhove, J., Tuerlinckx, F. & Lee, M. D. Hierarchical diffusion models for two-choice response times. Psychol. Methods 16, 44–62 (2011).

68. Wiecki, T. V., Sofer, I. & Frank, M. J. Hddm: Hierarchi cal bayesian estimation of the drift-diffusion model in python. Front. neuroinformatics 7, 14 (2013).

69. Ratcliff & Childers, R. Individual differences and fitting methods for the two-choice diffusion model of decision making. Decision 2, 237 (2015).

70. Spilcke-Liss, J., Zhu, J., Gluth, S., Spezio, M. & Glascher, J. Semantic incongruency interferes with endogenous attention in cross-modal integration of semantically congruent objects. Front. Integr. Neurosci. 13, 53 (2019).

71. Ratcliff, R., Philiastides, M. G. & Sajda, P. Quality of evidence for perceptual decision making is indexed by trial-to-trial variability of the eeg. Proc. Natl. Acad. Sci. 106, 6539–6544 (2009).

72. Philiastides, M. G., Heekeren, H. R. & Sajda, P. Human scalp potentials reflect a mixture of decision-related signals during perceptual choices. J. Neurosci. 34, 16877–16889 (2014).

73. Gamerman, D. & Lopes, H. F. (eds.) Markov chain monte carlo: stochastic simulation for bayesian inference (Taylor and Francis, London, 2006).

74. Gelman, A. & Rubin, D. B. Inference from iterative simulation using multiple sequences. Stat. Sci. 7, 457–472 (1992).

75. Spiegelhalter, D. J., Carlin, B. P. & Linde, A. V. D. Bayesian measures of model complexity and fit. J. Royal Stat. Soc. Ser. B (Statistical Methodol. 64, 583–639 (2002).

76. Etz, A. & Vandekerckhove, J. Introduction to bayesian inference for psychology. Psychon. Bull. & Rev. 25, 5–34 (2018).

77. Morey, R. D. & Rouder, J. N. Bayesfactor: Computation of bayes factors for common designs (r package version 0.9.12-4.2). retrieved from https://cran.r-project.org/package=bayesfactor.

78. Akiko Ikkai and Sangita Dandekar and Clayton E. Curtis. Lateralization in alpha-band oscillations predicts the locus and spatial distribution of attention. PLoS One 11, e0154796 (2016).

79. Palestro, J. J. et al.. A tutorial on joint models of neural and behavioral measures of cognition. J. Math. Psychol. 84, 20–48 (2018).

80. Bahg, G., Evans, D. G., Galdo, M. & Turner, B. M. Gaussian process linking functions for mind, brain, and behavior. Proc. Natl. Acad. Sci. 117, 29398–29406 (2020).

81. Turner, B. M., Wang, T. & Merkle, E. C. Factor analysis linking functions for simultaneously modeling neural and behavioral data. NeuroImage 153, 28–48 (2017).

82. Turner, B. M., Palestro, J. J., Miletić, S. & Forstmann, B. U. Advances in techniques for imposing reciprocity in brain-behavior relations. Neurosci. & Biobehav. Rev. 102, 327–336 (2019).

83. Turner, B. M., Rodriguez, C. A., Norcia, T. M., McClure, S. M. & Steyvers, M. Why more is better: Simultaneous modeling of eeg, fmri, and behavioral data. NeuroImage 128, 96–115 (2016).

